# Mobilization of tissue-resident memory CD4^+^ T lymphocytes and their contribution to a systemic secondary immune reaction

**DOI:** 10.1101/2020.04.02.021709

**Authors:** Carla Cendón, Weijie Du, Pawel Durek, Tobias Alexander, Lindsay Serene, Tina Lai, Axel Ronald Schulz, Anna Rao, Gitta-Anne Heinz, Ana-Luisa Stefanski, Anne Bruns, Katherina Siewert, Thomas Dörner, Hyun-Dong Chang, Hans-Dieter Volk, Chiara Romagnani, Mir-Farzin Mashreghi, Kevin Thurley, Andreas Radbruch, Jun Dong

## Abstract

While it is generally accepted that tissue-resident memory T lymphocytes protect host tissues from secondary immune challenges, it is unclear whether, and if so, how they contribute to systemic secondary immune responses. Here we show that in human individuals with an established immune memory to measles, mumps and rubella viruses, when challenged with the measles-mumps-rubella (MMR) vaccine again, tissue-resident memory CD4^+^ T cells are mobilized into the blood within 16 to 48 hours after vaccination. These cells then leave the blood again, and apparently contribute to the systemic secondary immune reaction, as is evident from the representation of mobilized T cell receptor Vβ clonotypes among newly generated circulating memory T lymphocytes, from day 7 onwards. Mobilization of the tissue-resident memory T cells is cognate, in that memory T lymphocytes recognizing other antigens, e.g. tetanus toxin, are not mobilized, unless they cross-react with the vaccine. These data originally demonstrate the essential contribution of tissue-resident memory T cells to secondary systemic immune responses, confirming that immunological memories to systemic pathogens are maintained (also) by tissue-resident memory T cells. In practical terms, the present work defines day 1 to 2 after antigenic challenge as a time window to assess the entire immunological T cell memory for a certain pathogen, including mobilized tissue-resident memory T cells, and its correlates of effectivity.

**Capsule summary:** The study demonstrates the rapid and cognate mobilization of tissue-resident memory CD4^+^ T cells into the blood upon antigenic rechallenge, and their contribution to secondary systemic immune responses.

## Introduction

Recent research has provided compelling evidence that, in addition to circulating central memory T cells (T_CM_) and effector memory T cells (T_EM_), which are readily detectable in the blood at all times [1], there are also significant populations of non-circulating, tissue-resident memory T lymphocytes. Tissue-resident memory T cells have been decribed for a variety of tissues, such as the skin, gut, lungs, liver [2–6], and bone marrow [7–11]. Compartments of tissue-resident memory T cells have been defined by confinement in parabiosis experiments [12, 13], by distinct gene expression signatures [4, 7, 11, 14–16], and by exclusive antigen receptor repertoires and specificities [3, 7, 11]. While tissue-resident memory T cells of epithelial tissue are capable of providing enhanced local protection at sites of previous infections [2, 3, 13, 17–20], tissue-resident memory T cells of bone marrow preferentially maintain long-term memory for systemic pathogens, like measles, mumps, and rubella viruses. In elderly individuals with high titers of measles specific antibodies, few if any measles-specific memory CD4^+^ T lymphocytes are detectable in the blood, but they are readily detectable in the bone marrow [7].

In the bone marrow, tissue-resident memory T cells reside individually in niches organized by stromal cells [11]. In both humans and mice, bone marrow memory CD4^+^ and CD8^+^ T cells maintain a non-proliferative, transcriptionally silent, resting state [7, 8, 11]. In secondary immune reactions, murine CD4^+^ tissue-resident memory T cells within the bone marrow form “immune clusters” and proliferate vigorously [21]. Whether or not tissue-resident memory CD4^+^ T cells, of the bone marrow or other tissues, are mobilized into the circulation in secondary immune reactions, and contribute to immune reactions in secondary immune organs, has been unclear so far.

Here we describe the mobilization of human CD4^+^ tissue-resident memory T cells into the blood, following a challenge with the live attenuated measles-mumps-rubella (MMR) vaccine. We show that MMR antigen-reactive CD4^+^ cells are rapidly mobilized into the blood within 16 to 48 hours after vaccination. They then disappear again from the blood, presumably participating in the secondary systemic immune reaction. Their T cell receptor repertoire becomes part of the repertoire of newly generated memory Th lymphocytes, demonstrating that tissue-resident memory T cells specific for systemic antigens can contribute essentially to systemic secondary immune responses.

## Methods

**Key resource** (see Supporting Information Table *S*1)

### Human Subjects

The recruitment of study subjects was conducted in accordance with the approval from the local ethical commission Charité-Universitätsmedizin Berlin (EA1/342/14) and informed consent in accordance with the Declaration of Helsinki. Fifteen healthy adults (5 male and 10 female; age±SEM = 34.67±2.11) with either documented measles, mumps, or rubella vaccination history or exposure through natural measles, mumps, or rubella infection were screened for their levels of pre-existing vaccine-reactive antibody titers (analyzed by Medizinisches Versorgungszentrum Labor Berlin). The donor’s previous antigen exposure either by vaccination or by natural infection was verified by detectable antigen-reactive antibody titers. This study discusses results obtained from 11 donors who received a live attenuated measles-mumps-rubella (MMR) (Priorix®, GSK, Germany) booster vaccination.

### Sample collection and preparation

Fifty millilitres of blood was drawn before vaccination (day 0), and at 6 time points (16 hours and days 1, 2, 3, 7 and 14) or at 2 time points (days 1and 14) after vaccination into Vacutainer® Heparin tubes (BD Biosciences, Plymouth, U.K.). Blood samples were subjected to immediate preparation and analyses. Mononuclear cells were isolated by density gradient centrifugation using Ficoll-Hypaque (Sigma-Aldrich). Serum samples were collected on days 0, 1, 3 and 14 before and after vaccination into Vacutainer® SST™ tubes (BD Biosciences, Plymouth, U.K.) and stored at −20°C until use.

### Flow cytometry analysis

Eight-to twelve-colour flow cytometry analysis was performed for the analysis of phenotype, cytokine profile, activation status, and cell sorting using a BD FACSAria cell sorter, LSRFortessa flow cytometer (BD Biosciences), or MACSQuant (Miltenyi Biotec). The following anti-human antibodies were used to stain cells: CD3, CD19, CD14, CD4, CD8, CD45RA, CD25, CD127, CCR7, Ki67, CD154, CD69, IFNγ, IL-2, TNFα, and fixable Live/Dead or propidium iodide (PI). Stained cells were acquired using FACSDiva (BD Biosciences) or MACSQuantify™ software and data were analyzed with FlowJo (Tree Star).

### Quantification of leukocytes

A 50 μL aliquot of each whole blood sample was stained immediately after drawing with 50 μL of a fluorochrome-conjugated mouse anti-human antibody mixture (CD45, CD56, CD19, CD14, CD3, CD4 and CD8) in the presence of FcR-blocking reagent (1:5). After staining, erythrocytes were lysed by adding 500 μL of Buffer EL and incubating for 30 min at 4°C. A defined volum of each sample was acquired on a MACSQuant (Miltenyi Biotec). The resulting leukocyte population cell counts were used as the basis for calculating subpopulations.

### Identification of antigen-reactive memory CD4^+^T cells

PBMCs were cultured in RPMI 1640 supplemented with 1% GlutaMAX, 100 U/mL penicillin, 100 μg/mL streptomycin, and 5% (vol/vol) human AB serum. Anti-CD28 (1 μg/mL), measles (5μg/mL), mumps (5μg/mL), rubella (10 μg/mL) antigen, and tetanus toxoid (TT) [1 lethal facor (Lf/)mL)] were used to stimulate cells. Stimulation with CD28 alone and in combination with Staphylococcal enterotoxin B (1 μg/mL) were used as negative and positive controls, respectively. For each condition, at least 5-10 × 10^6^ cells were stimulated for 7 h at 37° and 5% CO_2_, in the presence of brefeldin A (1 μg/mL) for the last 4 h. Stimulated cells were first stained for surface antigens followed by fixation with BD FACS Lysing and Perm2 solutions according to the manufacture’s instruction. The antigen-reactive memory CD4^+^ T cells were identified as Live/Dead^−^CD19^−^CD14^−^CD3^+^CD8^−^ CD4^+^CD45RA^−^CD154^+^cytokine^+^ and the proliferating antigen-reactive cells as Live/Dead^−^CD19^−^ CD14^−^CD3^+^CD8^−^CD4^+^CD45RA^−^CD154^+^Ki67^+^. At least 10^6^ lymphocytes were analyzed. Alternatively, antigen-reactive CD154^+^ memory CD4^+^ T cells were isolated using the CD154 MicroBead Kit. In brief, cells were indirectly magnetically labeled with CD154-biotin and anti-biotin Microbeads and enriched by two sequential MS MACS columns. Cell surface staining was performed on the first column, followed by fixation, permeabilization, and intracellular cytokine staining on the second column using a established protocol [22].

### Isolation of antigen-reactive memory CD4^+^ T cells

At least 5 × 10^6^ PBMCs were stimulated for 7 h with antigen in the presence of 1 μg/mL anti-CD40. Stimulated cells were stained for 15 min at room temperature with the following antibodies: CD3, CD4, CD8, CD19, CD14, CD45RA, CD69 and CD154. Antigen-reactive memory CD4^+^ T cells were identified by the expression of PI^−^CD19^−^CD14^−^CD3^+^CD8^−^CD4^+^CD45RA^−^CD154^+^CD69^+^. Measles-reactive memory CD4^+^ T cells sorted before, days 1 and 14 after vaccination were further processed for TCR CDR3 Vβ sequencing. In addition, measles- and TT-reactive memory CD4^+^ T cells sorted 1 day after vaccination were used for the generation of antigen-reactive T cell lines.

### Expansion and re-stimulation of antigen-reactive memory CD4^+^ T cell lines

Purified antigen-reactive CD69^+^CD154^+^ memory CD4^+^ T cells (as described earlier) were cultured with CD3 depleted and irradiated (40 Gy on ice for 35 min) autologous feeder cells at ratios of 1:100 in a 48-well plate in X-VIVO15 medium supplemented with 100 U/mL penicillin, 100 μg/mL streptomycin, 5% (vol/vol) human AB serum, and 200 IU/mL IL-2. Cells were expanded for 14 days, with 200 IU/mL IL-2 supplemented every 3 days during the first 10 days of the cell culture.

For re-stimulation, freshly isolated autologous PBMCs were labelled with CFSE (1 μM) following the manufacturer’s protocol. CFSE labeled PBMCs were co-cultured with 2,5 × 10^5^ expanded T cells at ratios of 2:1 and stimulated with different antigens (measles, mumps, rubella, TT, or CMV) in the presence of CD28 (1 μg/mL) for 6 h, with the last 4 h with brefeldin A (1 μg/mL). Stimulated cells were stained with surface antigens and intracellular cytokines.

### Determination of antigen-specific antibodies by Enzyme-linked Immunosorbent Assay (ELISA)

Before and after MMR vaccination, antigen specific IgM and IgG responses against measles, mumps, or rubella, and the IgG response against TT in the serum samples were analyzed with the antigen-specific IgM and/or IgG ELISA following the manufacturer’s instruction.

### TCR CDR3 Vβ gene library preparation

Total RNA was extracted from 2500 measles-reactive memory CD4^+^ T cells using the Qiazol Lysis Reagent and a miRNAeasy Micro Kit, following the manufacturer’s protocols. The resulting RNA concentration and quality were determined in a Bioanalyzer. Eight μL RNA were then processed for the TCR CDR3 Vβ library preparation, following a published protocol [23] with modifications. Briefly, the first strand of complementary DNA (cDNA) with the 5’ template switch adapter SmartNNNa was synthesized using a SMARTScribe reverse transcriptase, following the manufacturer’s instruction. cDNAs were purified using a MinElute PCR purification Kit prior to a two-step PCR with a proof-reading Q5 DNA polymerase and barcoded primers (see key resources table). PCR products of each step were purified using a QIAquick PCR purification kit, following manufacturer’s instruction. The second round of PCR products containing fragments of about 550 bp were sliced under a Blue LED Transilluminator (Herolab, Germany) and further purified by PCR-clean-up Gel extraction and stored at −20°C until adaptor ligation and sequencing. The concentration of the size-selected PCR products was determined using a Qubit fluorometer. Finally all barcoded PCR products/samples were mixed together at equal ratios, and the Illumina adapters were ligated according to the manufacturer’s protocol. The pooled libraries were purified by PCR-clean-up Gel extraction.

### TCR CDR3 Vβ gene sequencing and data analysis

Paired-end (2 × 150bp) sequencing of the TCR CDR3 Vβ gene transcripts was performed on an Illumina NextSeq 500 platform. Inline barcodes were demultiplexed and the Illumnia Adapters removed by Illuminas bcl2fastq 1.8.4 software. De-multiplexing of sample barcodes as well as the extraction of the CDR3 Vβ gene sequences were performed by the open source software tools migec 1.2.4 [24] and VDJtools [24] using the unique molecular identifier (UMI) guided assembling and error correction with default setting, minimal number of 5 reads per UMI and blast. Data of biological replicates at the same time point were either pooled or merged for overlap if specified. For downstream analysis and plotting, base-level R functions and the public available software package VDJtools[25] were used. The inverse Simpson index is a measure of diversity commonly applied in ecology, it can be computed as 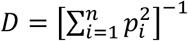, where the *p*_*i*_ are relative frequencies of all *n* observed clonotypes. If all clonotypes have equal frequency, i.e. *p*_*i*_ = 1/*n* for all *i*, then *D* = *n*, and therefore *D* can be interpreted as the “effective number of parties” generating the observed diversity in a given sample. Although the inverse Simpson index is a comparatively robust measure of diversity [26], for direct comparisons between experiments we adjusted the total number of reads by “down-sampling”, i.e. random selection of samples until the number of reads in the smallest sample is reached.

### Quantification and statistical analyses

Polyfunctionality of antigen-reactive memory CD4^+^ T cells was analyzed by using Flowjo Boolean (logic) operators (AND and NOT) of overlapping regions to define all possible combinations of cytokine profiles. The results were displayed as an absolute number of cytokine-producing cells per 10^6^ CD4^+^ T cells or per 10^3^ CD154 MACS enriched cells.

Statistical analysis for differences in antigen-reactive memory CD4^+^ T cells induced by MMR vaccination were performed by one-tailed paired *t*-test to test for an early increase of T cell numbers in individual donors.

Comparison of intersections of TCR repertoire samples after repetitive down-sampling was performed using a rank-based permuation test (method “oneway” from the R-based “coin” package).

In all analyses, *p*-values ≤ 0.05 were considered to be statistically significant.

### Data and Code Availability

The TCR CDR3 Vβ gene sequencing datasets generated during this study will be deposited in the Gene Expression Omnibus (GEO) database, www.ncbi.nih.gov/geo.

## Results

### Mobilization of memory CD4^+^ T cells into the blood in a secondary immune response

Eleven healthy human volunteers (MMR1-11), aged 28 to 43 years, were immunized subcutaneously with the live attenuated measles-mumps-rubella virus vaccine, as outlined in Fig 1A. Prior to vaccination, serum IgG titers, specific for each of the three viruses, had been determined. Except vaccinee MMR4, who lacked antibodies specific for measles virus, all vaccinees had pre-existing antibodies specific for each of the three viruses, indicating that they had an immunological memory to the viruses, prior to vaccination (Table S2). Virus-reactive memory CD4^+^ T lymphocytes were identified upon restimulation with antigen *ex vivo* as cells expressing CD154 (CD40L) and one or more of the cytokines IL-2, TNF-α, and IFN-γ, as described previously [7, 22] (Fig S1, *A* and *B*), at time points indicated in Fig 1A. Frequencies of antigen-reactive Th cells were highly reproducible between measurements for technical replicates (Fig S2, *A*) as well as for biological replicates, i.e. blood samples of individual donors, taken at two consecutive days without vaccination (Fig S2, *B*).

**Figure 1.**
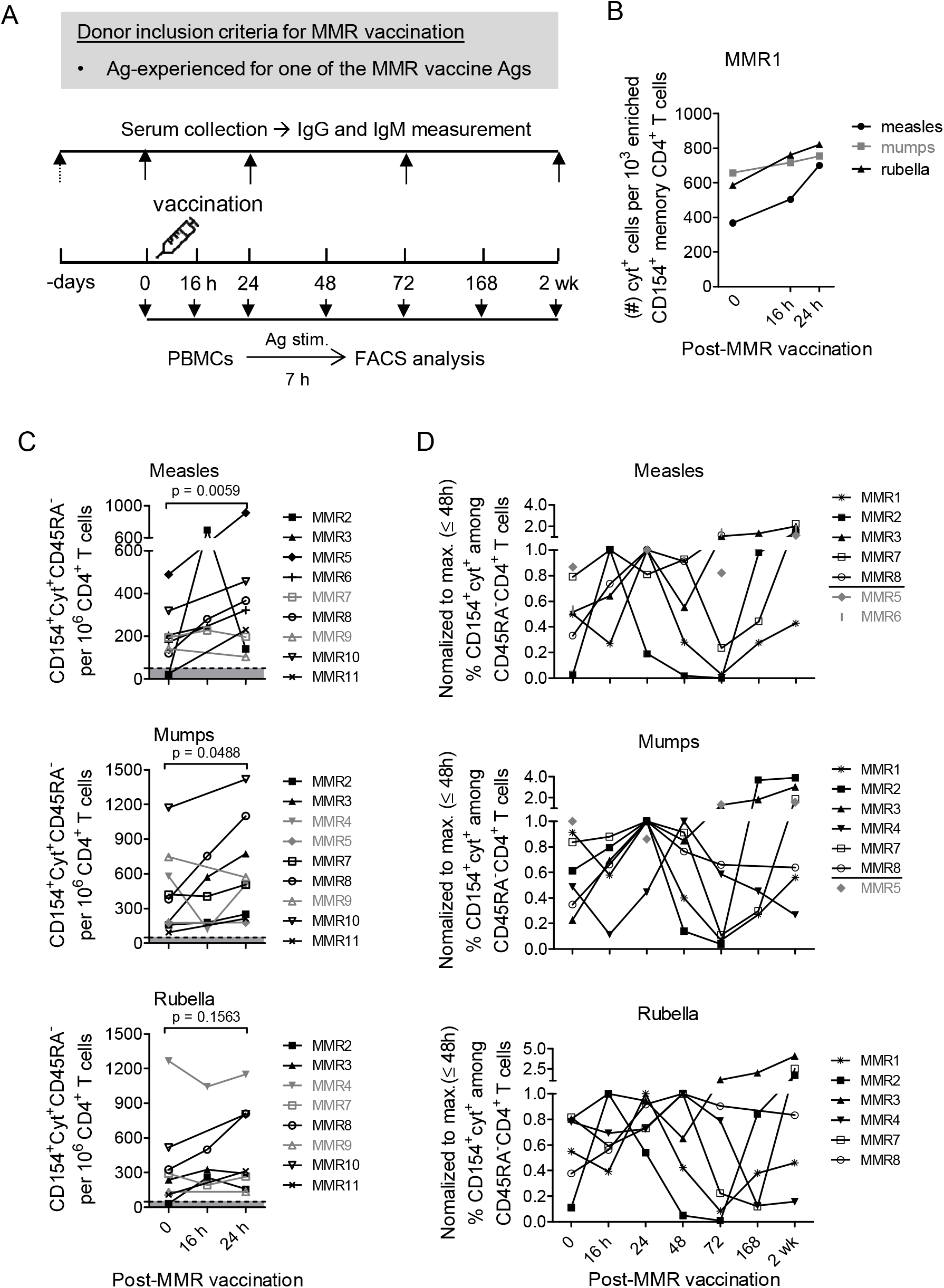
Kinetics of virus-specific memory CD4^+^ T cell responses after MMR vaccination. (A) Donor inclusion criteria and scheme for analyzing virus-specific memory CD4^+^ T cell responses following MMR booster vaccination. PBMCs stimulated with measles, mumps, or rubella and analyzed for antigen-reactive CD154^+^cytokine^+^ memory T cells (memory CD4^+^ T cells expressing CD154 (CD40L) and one or more of the cytokines IL-2, TNF-α, and IFN-γ) per 10^3^ enriched CD154+ memory CD4^+^ T cells (B) or per 10^6^ CD4^+^ T cells (C) and in relative frequencies to its maximum by 48 hours after vaccination (D), at indicated time points. In C, the dotted black line indicates the minimum threshold for detection with the assay and the grey area underneath values below reliable detection limit. Significance was determined by paired t-test with Welch’s correction and one-tailed *P* value was shown. In both B and C, data shown in grey indicate the responses at 24-hour that are not more than 30% greater than before vaccination. In D, data shown in grey indicate vaccinees analyzed on the following time points: before (0), 24-, 72-hour, and 2wk after vaccination.

For the 11 vaccinees immunized with the MMR vaccine, we monitored the frequencies of virus-reactive memory Th cells over time in 11 of them for measles, 10 for mumps, and 9 for rubella. Strikingly, vaccine-reactive memory Th cells were rapidly mobilized into the blood in most, though not all vaccinees, leading to a significant numerical increase in memory CD4^+^ T cells reacting to measles and mumps, within 16 to 24 hours after vaccination (Fig 1, *B* and *C*). By 24 hours post-vaccination, differences in frequencies of measles-, mumps-, and rubella-reactive memory CD4^+^ T cells in the blood increased significantly compared to time-matched “baseline” control experiments without vaccination (Fig S3), and in 20 of 29 memory T cell responses analyzed by more than 20% (Fig 1, *B* and *C*). Twenty of the 29 memory immune reactions were also analyzed at 48- and/or 72-hour after vaccination (Fig S4, *A*). In 17 of these 20 memory immune reactions, virus-specific CD154^+^cytokine^+^ memory CD4^+^ T cells peaked at 16-, 24-, or 48-hour after vaccination and thereafter dropped to their baseline levels or below (Fig 1, *D*). Taken together, vaccination of previously immune donors rapidly mobilizes antigen-reactive Th cells from the tissues into the blood, within 1 to 2 days, and these cells disappear again from the blood on days 3 and 4.

Fourteen days after vaccination, antigen-reactive memory T cells circulating in the blood increased significantly (Fig S4, *B*). In 26 of the 30 immune reactions monitored (Fig S4, *A*), 30% more virus-reactive memory T cells were detected, as compared to d0, before vaccination (Fig S4, *B*). It is interesting to note that only 13 out of 24 memory T cell-generating immune responses analyzed also showed an increase in virus-specific serum IgG antibody titers (Fig S4, *C*). Also interesting is that the magnitude of memory T cell responses 2 weeks after vaccination did not significantly correlate with specific antibody titers before vaccination (Fig S5).

### Rapidly mobilized CD4^+^ memory T cells are activated, non-proliferative and polyreactive

In accordance with the rapid mobilization of vaccine-specific memory T cells into the blood, as defined by their reactivation *ex vivo*, we detected a rapid mobilization, at 16 to 48 hours, of *in vivo* activated memory T cells in four vaccinees analyzed, MMR2, MMR3, MMR7 and MMR8. Activation of these cells *in vivo* was evident from their expression of CD154 (Fig 2, *A*) or any of the cytokines IL-2, IFN-γ or TNF-α (Fig 2, *B*) determined directly *ex vivo*, without antigenic restimulation. Since their appearance parallels that of vaccine-reactive Th cells, it is tempting to speculate that they had been activated by the vaccine *in vivo*.

**Figure 2.**
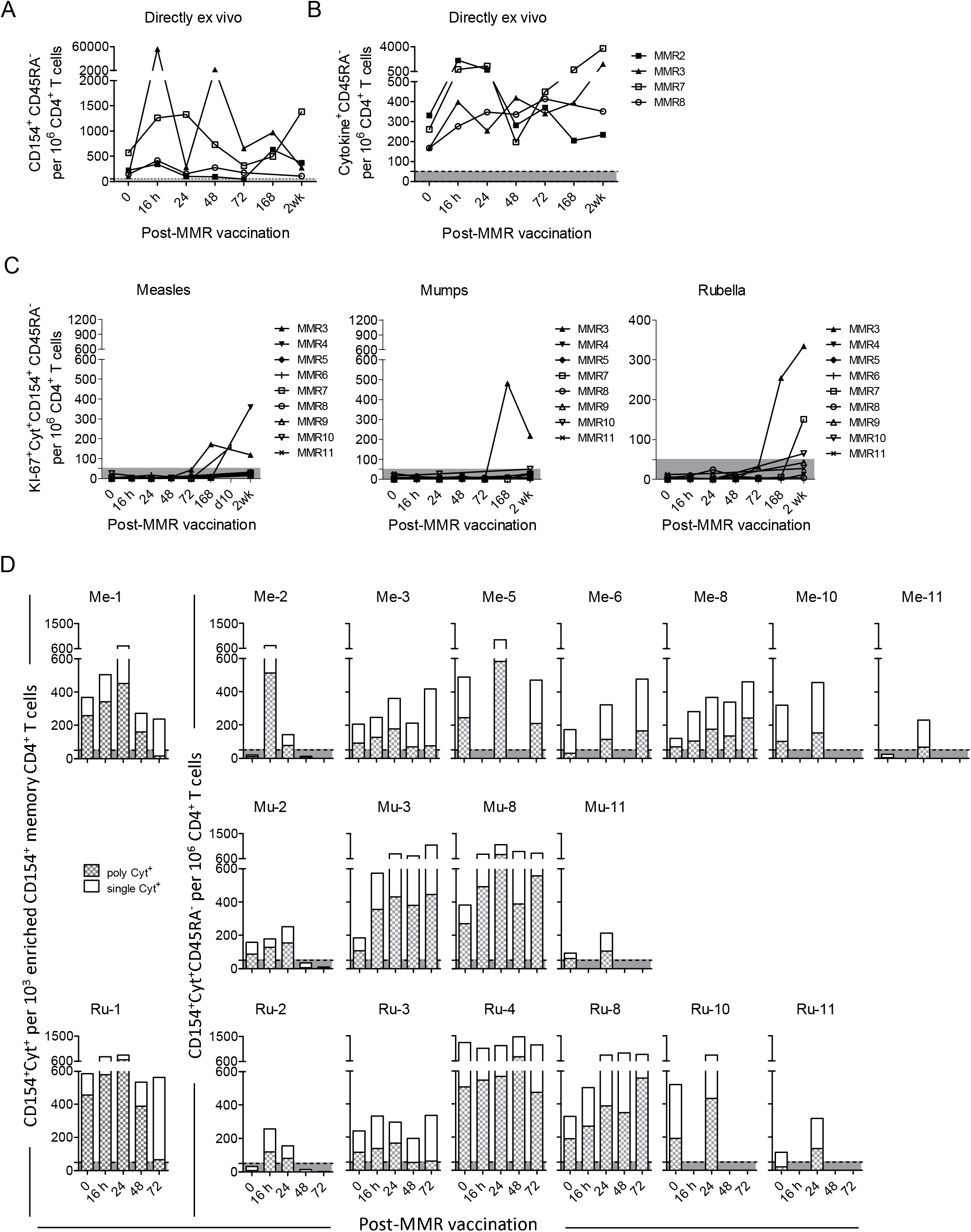
Characterization of memory CD4^+^ T cells following MMR vaccination. (A) PBMCs from vaccinees analyzed, directly *ex vivo*, for CD154^+^ (A) and CD154^+^cytokine^+^ (B) CD45RA^−^ memory T cells per 10^6^ CD4^+^ T cells before and at indicated time points after vaccination. (C, D) PBMCs from vaccinees stimulated with measles, mumps, or rubella and analyzed for antigen-reactive Ki-67^+^CD154^+^cytokine^+^ memory T cells (C) and single, double, or triple cytokine-producing subpopulations of CD154^+^cytokine^+^ memory T cells (D) per 10^6^ CD4^+^ T cells before and at indicated time points after vaccination. In C and D, the dotted black line marks the assays’s minimum threshold for detection, and the grey areas underneath, values below reliable detection limit.

It is a matter of debate whether memory T cells, tissue-resident or circulating, are maintained as non-proliferative, resting cells. Here we show that until 72 hours after vaccination, the numbers of Ki-67 expressing virus-reactive memory T cells in the blood of the vaccinees were below the level of detection (Fig 2, *C*; S6), implying that the mobilized memory T cells are not expressing Ki-67. Later, on d7 and d14 after re-immunization, antigen-reactive Ki-67^+^CD154^+^cytokine^+^ memory CD4^+^ T cells were readily detectable in the blood (Fig 2, *C*; S6).

Tissue-resident memory T cells, in particular the measles-reactive ones of the bone marrow, in contrast to their circulating counterparts, have been shown to express several cytokines simultaneously upon reactivation [7]. Here we show that upon MMR revaccination, in 19 out of 29 memory T cell responses with mobilized memory CD4^+^ T cells, the majority of the CD4^+^ T cells, peaking on day 1 express several cytokines simultaneously, i.e. they are “polyfunctional” (Fig 2, *D*; S7; S8) [27].

### Mobilization of memory Th cells by MMR vaccine is cognate

The rapid mobilization of vaccine-reactive memory Th cells into the blood was not due to a general increase in numbers of peripheral T cells and memory T cells (Fig S9). To determine whether the rapid mobilization of vaccine-reactive memory T cells is cognate or not, we determined whether the MMR vaccine also mobilized memory T cells recognizing the antigen tetanus toxoid (TT). Surprisingly, we observed a rapid mobilization of TT-reactive memory T cells in 3 out of 4 vaccinees analyzed (Fig 3, *A*). Moreover, these vaccinees also showed a drop in numbers of TT-reactive memory T cells on d2 and/or d3, and their reappearance on d7/14 (Fig 3, *A*), resembling the kinetics of vaccine-reactive memory T cells (Fig S4, *A*).

**Figure 3.**
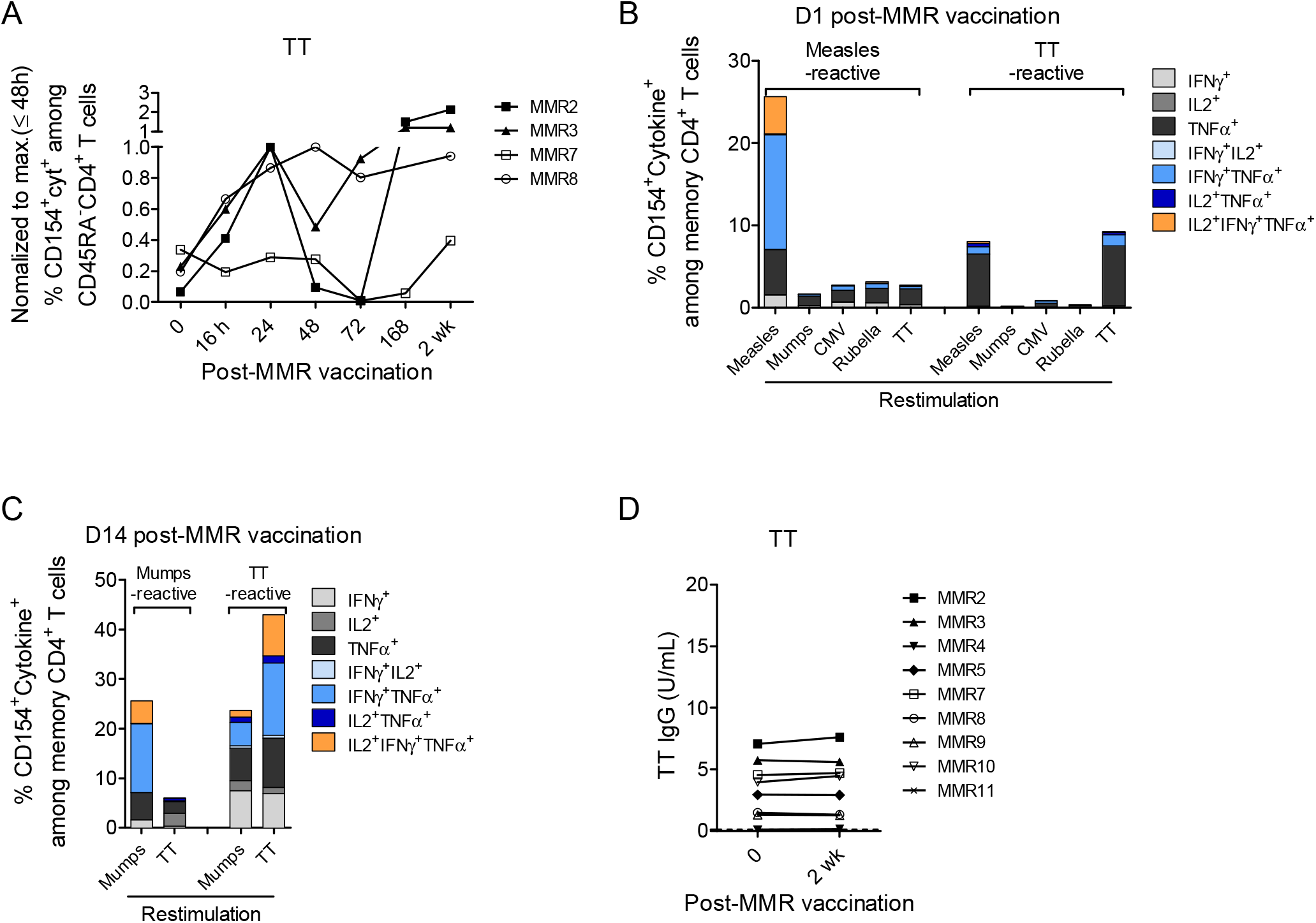
Specificity of antigen-reactive memory CD4^+^ T cell responses to MMR vaccine. (A) Mobilization of TT-reactive memory T cells following MMR booster vaccination. PBMCs from four vaccinees stimulated with TT and analyzed for TT-reactive CD154^+^cytokine^+^ CD45RA^−^ memory CD4^+^ T cells. (B, C) Specificity of expanded measles- or TT-reactive CD154^+^ memory CD4^+^ T cell lines isolated after MMR vaccination. PBMCs from MMR vaccinees MMR8 (B) and MMR7 (C) were stimulated with measles or TT 1 (d1) or 14 (d14) days after MMR vaccination. Antigen-reactive CD154^+^ memory CD4^+^ T cells were then isolated from these samples and expanded for 14 days with IL-2 and autologous APCs. Expanded cell lines were restimulated with the indicated antigens in the presence of autologous APCs and the reactive cells were identified by coexpression of CD154 and one or more of the cytokines IL-2, TNF-α, and IFN-γ. (D) The production of IgG antibodies specific for TT antigen before and after MMR vaccination. The dotted black line marks the assay’s minimum threshold for detection, and the grey area underneath, values below reliable detection limit.

To investigate whether the mobilized TT-reactive memory T cells were crossreactive to the MMR vaccine, TT-reactive memory T cells of vaccinee MMR8 were isolated on d1 after MMR vaccination, based on co-expression of CD154 and CD69. These cells were then expanded for 2 weeks with IL-2, then restimulated with measles, mumps, rubella, CMV or TT. TT-reactive T cell lines isolated after vaccination were reactive to both TT and measles, but not to mumps or rubella (Fig 3, *B*), suggesting that cross-reactive memory T cells had been mobilized in this vaccinee, and that mobilization indeed had been antigen-specific. We also detected mobilized crossreactive memory T cells on later time points after MMR vaccination. In that case, we isolated TT-reactive cells from vaccinee MMR7 on d14 after vaccination, expanded the isolated cells, and restimulated with mumps antigen (Fig 3, *C*). Even though crossreactive TT-reactive memory T cells were mobilized, and apparently also activated by MMR vaccination, this did not have a detectable impact on the TT-specific serum antibody titers (Fig 3, *D*).

### Rapidly mobilized memory Th cells contribute to the systemic secondary immune reaction

To determine whether the mobilized vaccine-reactive memory T cells contribute to the immune response triggered by the vaccine, we compared the T cell receptor (TCR) CDR3 Vβ clonotypes of the rapidly mobilized memory T cells to those of the memory T cells present in the blood 14 days after vaccination. To this end, two replicates with equal numbers (2.5 × 10^3^) of measles-reactive CD154^+^CD69^+^ memory CD4^+^ T cells were isolated from 30 mL of peripheral blood drawn from each of the vaccinees MMR10 and MMR11 before (d0), and 1 (d1) and 14 (d14) days after MMR vaccination. The global CDR3 Vβ clonal repertoires of measles-reactive memory CD4^+^ T cells at these three time points were then compared (Table S3; Fig 4, *A*; Fig S10, *A*).

**Figure 4.**
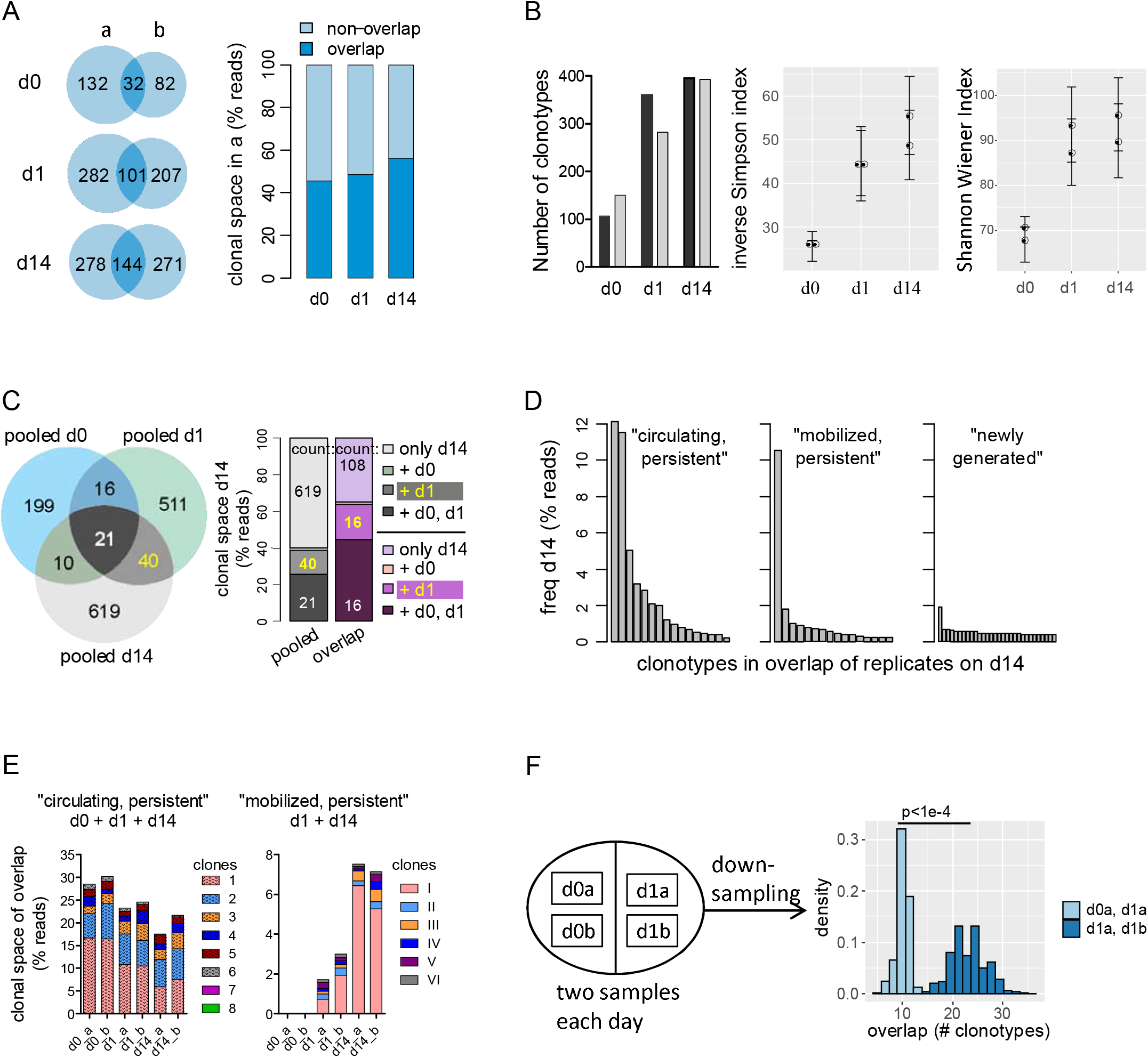
TCR CDR3 Vβ clonotypes analysis of measles-reactive CD4^+^ memory T cells after MMR vaccination. PBMCs were isolated before (d0) and 1 (d1) and 14 (d14) days after MMR vaccination from vaccinee MMR10, restimulated *ex vivo* with measles for measles-reactive CD154^+^CD69^+^ memory CD4^+^ T cells. Analogous data for vaccinee MMR11 are shown in Fig. S10. (A) Analysis of two replicates at each time point, shown is the “overlap”, i.e. the amount of clonotypes detected in both replicates, in terms of clonotype count (Venn diagrams) and frequency (bar graphs). (B) Diversity of the TCR repertoire. Shown are the number of clonotypes, the inverse Simpson index and the Shannon Wiener Index at three time points, for both replicates in A. Error bars indicate standard deviation after repetitive down-sampling. (C) Venn diagrams show comparative numbers of clonotypes in “pooled” repertoires, i.e. clonotypes detected in at least one replicate. Clonal space (bar graph) corresponds to the fraction of reads detected on d14, “overlap” or “pooled”, that were additionally present in at least one of the replicates at indicated time points. Numbers in the bar plot indicate numbers of clonotypes (for pooled replicates, these numbers match those of the Venn diagram). (D) Frequency distributions of “circulating, persistent” (d14+d0+d1), “mobilized, persistent” (d14+d1) and “new” (only d14) clonotypes (see C), with respect to the overlap of replicates on d14. For “new”, only the 30 most frequent clonotypes are shown. (E) Contribution of each replicate to the “circulating, persistent” and “mobilized, persistent” clonotypes. Clonal space is given as percentage of reads in the overlap of replicates on indicated days, and only clonotypes present in all relevant samples are shown. (F) Comparison of the overlap of samples taken on different days or taken on the same day. Shown are the distributions of indicated sample overlaps after repetitive down-sampling to the number of reads in the smallest among all samples taken on d0 and d1 (see also Fig. S10, *F*).

The number of CDR3 Vβ clonotypes increased by ≥75% from d0 to d1 in both vaccinees, MMR10 (Fig 4, *B*) and MMR11 (Fig S10, *B*) and across all measured samples. The clonal diversity increased from d0 to d1 in both donors as well, as quantified by the inverse Simpson index, a measure of diversity that can be interpreted as the “effective number of parties” in a given sample [26] (Fig 4, *B*; Fig S10, *B*). By d14, the diversity and number of clonotypes stayed high in MMR10 (Fig 4, *A*), and decreased to basal levels in MMR11 (Fig S10, *A*), in line with their T cell response kinetics (Fig S4, *A*). Another measure of diversity, the Shannon Wiener Index (Fig 4, *B*; Fig S10, *B*) gave qualitatively similar results. Together, these results demonstrate that MMR vaccine triggers mobilization of antigen-reactive memory CD4^+^ T cells expressing distinct clonotypes, which had been absent from the repertoire of pre-existing circulating memory T cells.

To assess the contribution of the mobilized T cells to the repertoire on d14, we analyzed the clonal space at this time in more detail. To this end, we considered both the “pooled” clonal space (clonotypes detected in either of the replicates) and the “overlap” clonal space (clonotypes detected in both replicates) (Fig 4, *A* and *C*; Fig S10, *A* and *C*). In both vaccinees, the largest fraction of antigen-reactive CD4^+^ T cells on d14 consisted of clonotypes that had not been detected in the blood on either d0 or d1, i.e. 619 clonotypes for MMR10 and 488 clonotypes for MMR11, comprising 60% of the pooled clonal space for MMR10 and 80% for MMR11 (Fig 4, *C*). Twenty-one clonotypes of MMR10 and 22 of MMR11, about 25% and 10% of the pooled clonal space on d14, respectively, were “circulating” clonotypes, detected on d0, d1 and d14. Fourty clonotypes of MMR10 and 60 of MMR11, comprising about 15% and 10% of the pooled clonal space on d14, respectively, were “mobilized” clonotypes, i.e. detected on d1 and d14, but not on d0 (Fig 4, *C*; Fig S10, C; highlighted in yellow). In the “overlap” clonal space, representing clonotypes detected in both replicates of each day from a given vaccinee, the “mobilized” clonotypes, i.e. those found on d1 but not on d0, covered about 20% for MMR10 and 10% for MMR11 of the repertoire on d14.T he “circulating” clonotypes covered about 40% and 15% of the repertoire on d14, of MMR10 and MMR11, respectively (Fig 4, *C*; Fig S10, *C*). Remarkably, we detected 3 highly abundant clonotypes with more than 5% of reads in donor MMR10, one of them “mobilized”, two “circulating” (Fig. 4, D). In MMR11, no highly abundant clones were detected on d14 (Fig. S10, *D*), in agreement with the reduced Simpson diversity in this case (Fig. S10, *B*). In both MMR10 and MMR11, individual “circulating” and “mobilized” clones were consistently detectable in each of the replicates, maintained their clonal space or even expanded, such as clone I from MMR10 (Fig 4, *E*; Fig S10, *E*).

To exclude the option that detection of increased diversity on d1, as compared to d0, was merely due to the higher sampling rate on d1 (Table S3), we tested the “null hypothesis” that the repertoires on d0 and d1 essentially had been the same. If that had been the case, and all samples taken on d0 and d1 were representing the same repertoire, only at varying sampling effectivity, the overlaps between replicates of each day would be the same as those between replicates of different days. However, this is not the case: after repetitive random downsampling of the number of reads to those of the smallest sample, the distribution of overlaps between replicates of the same day was significantly different from the distribution of overlaps between replicates representing different days (Fig. 4, *F*; Bonferroni corrected *p*-values < 10^−4^). This was true for all possible combinations of samples in both donors, ruling out the null hypothesis (Fig. S10, *F*), and demonstrating that indeed the repertoires of d0 and d1 were different, because of the mobilization of memory T cells with additional clonotypes.

Taken together, measles-specific CD4^+^ T cell clones mobilized into the blood by reimmunization with the MMR vaccine contribute significantly to the systemic secondary immune reactions against measles.

## Discussion

The discovery of memory T lymphocytes residing in distinct tissues, like the skin [2], lungs [4, 13] and bone marrow [7, 11] has revealed a considerable compartmentalization of immunological memory. Tissue-resident memory T lymphocytes are generally considered to protect their tissue of residence directly against repeated antigenic challenges [13, 18, 28]. Their contribution, as compared to circulating memory T lymphocytes, to systemic secondary challenges has been less clear, although the pioneering work of McGregor and Gowans has demonstrated, more than 50 years ago, that secondary immune reactions are not impaired by ablation of circulating lymphocytes [29].

In the present study we have tracked CD4^+^ memory T lymphocytes specific for the viruses measles, mumps or rubella, in the peripheral blood of human adult donors immunized with a combined measles-mumps-rubella (MMR) vaccine, over two weeks. All but one of the donors had pre-existing immunological memory for all of these viruses, i.e. virus-specific IgG antibodies. Different from our previous cohort of donors aged 40-70 years, most of whom did not have detectable numbers of measles-reactive memory Th cells in the blood [7]; many of the present study subjects aged 28-43 years had detectable virus-reactive memory Th cells in the blood, before vaccination (d0). Irrespective of this, in 20 of 29 secondary immune responses analyzed, we observed a rapid mobilization of virus-reactive memory CD4^+^ T cells into the blood, between 16 and 24 hours after immunization. Such cells then disappeared again from the blood, between d2 and d3, and appearred again in the blood from d7 on, with their numbers increasing until d14, the last point of analysis. The rapid mobilization of CD4^+^ T cells into the blood is antigen-specific (cognate), and contributes to the systemic immune response, as is evident by the presence of their antigen-receptor clonotypes among those of the memory T cells isolated from blood on d14.

Antigen-specific memory provided by the adaptive immune system is multi-layered. Antibodies secreted by memory plasma cells provide immediate protection, while memory T and B lymphocytes provide reactive memory against excessive or modified antigen [9, 30]. Moreover, immune memory cells are compartmentalized in various tissues [9, 31]. We have previously shown that prominent populations of memory plasma cells and memory CD4^+^ and CD8^+^ T lymphocytes reside and rest in the bone marrow [7, 8, 11, 16]. Memory CD4^+^ T lymphocytes of the bone marrow store longterm memory to systemic pathogens, like measles, mumps and rubella, even in the apparent absence of such memory cells from the blood [7]. Tissue-resident memory CD4^+^ T cells are heterogeneous, with some of them expressing CD69 and others not, and an ongoing debate on the transcriptional identity and lifestyle of tissue-resident memory CD4^+^ T cells [7, 9, 10, 14, 16, 32]. Here we demonstrate the rapid mobilization of CD4^+^ T cells specific for measles, mumps or rubella into the blood upon antigenic challenge, and their contribution to the systemic immune response. Since they obviously have been activated and mobilized, we cannot conclude on their tissue-of-origin, nor their signature transcriptome as resting cells, other than stating that they had been resident before vaccination, and that they are polyfunctional in terms of cytokine expression. Polyfunctionality is a decisive property of MMR-specific bone marrow resident memory CD4^+^ T cells [7].

Ten of the 11 immunized vaccinees (MMR1-3 and MMR5-11) had pre-existing memory to all three viruses. Only one donor, MMR4 did not have measles-specific IgG antibodies on d0. The 29 individual memory immune reactions of 11 donors analyzed displayed a considerable heterogeneity. For example, the rapid mobilization of virus-specific tissue-resident memory CD4^+^ T lymphocytes was independent of pre-existing antibody titers, e.g. MMR6 vs measles, and MMR8 vs measles. Mobilization of virus-specific tissue-resident memory CD4^+^ T cells also did not correlate to numbers of pre-existing circulating memory cells of the same specificity, e.g. MMR11 vs mumps, and MMR5 vs mumps. Moverover, mobilization of virus-specific tissue-resident memory CD4^+^ T cells did not correlate to efficacy of the subsequent immune reaction in generating new specific antibodies (e.g. MMR2 vs rubella, and MMR3 vs rubella) or increased numbers of circulating memory T cells of that specificity on day 14 (e.g. MMR2 vs measles, and MMR8 vs measles). Thus the mobilization of virus-specific tissue-resident memory CD4^+^ T cells, triggered by vaccination, is an independent event, and presumably just reflects the availability of tissue-resident memory CD4^+^ T cells in a given vaccinee.

Although the molecular details of mobilization are not clear, nor is the tissue from which the cells are mobilized, we can show here that the mobilization is cognate, i.e. only antigen-reactive CD4^+^ T cells are mobilized. In the vaccinees analyzed, the overall numbers of memory CD4^+^ T lymphocytes in the blood did not change significantly after vaccination. Although we observed mobilization of memory CD4^+^ T cells recognizing tetanus toxoid (TT), an antigen not present in the MMR vaccine, only TT-reactive CD4^+^ T cells crossractive to the vaccine were mobilized, in line with the notion that mobilization is cognate. This cognate mobilization of tissue-resident memory CD4^+^ T cells requires antigen-presenting cells to scan tissue(s) for reactive CD4^+^ T cells, and activate them to egress into the blood within 1 day after re-immunization.

A fast kinetic of reactivation of tissue-resident memory T cells is also observed within their host tissue. As we have previously shown, in the bone marrow, memory CD4^+^ T lymphocytes individually localize to stromal cells, and MHC class II-expressing antigen presenting cells are not in their vicinity [11, 21]. However, upon cognate reactivation *in situ*, murine bone marrow memory CD4^+^ T cells are rapidly mobilized from their niches, aggregate in “immune clusters” with antigen-presenting cells and proliferate vigerously [21]. In the present study, the mobilized tissue-resident memory CD4^+^ T cells did not yet proliferate on d1, suggesting that they had left their tissue niches and directly egressed into the blood.

In functional terms, the mobilized memory CD4^+^ T cells did contribute significantly to the systemic secondary immune reactions. In the vaccinees analyzed, T cell receptor clonotypes of the mobilized memory CD4^+^ T cells, i.e. those cells detected on d1 but not on d0, were also present, some even expanded, among memory CD4^+^ T cells of d14. In the relatively young population of vaccinees analyzed here, 28 to 43 years of age, circulating and mobilized tissue-resident memory CD4^+^ T cells contributed about equally to the systemic secondary immune reaction. We had shown before, from a cohort of subjects at 40-70 years of age, that circulating memory CD4^+^ T cells disappear, while tissue-resident memory CD4 T cells specific for measles, mumps and rubella remain easily detectable in the bone marrow [7]. The present results would predict that in aged humans the systemic secondary immune response to childhood pathogens is entirely dependent on mobilized tissue-resident memory CD4^+^ T cells. Those are well equipped to protect, though, since they are polyfunctional, i.e. can express several cytokines orchestrating protective anti-viral immune responses [33].

The key observation of cognate mobilization into the blood of CD4^+^ tissue-resident memory T cells, reactive to systemic pathogens like measles, mumps and rubella, including memory CD4^+^ T cells crossreactive to entirely differenct antigens, in secondary immune reactions, and their participation in the systemic immune reaction to those viruses, demonstrates that different compartments of immunological memory cooperate to provide efficient systemic immunity. Beyond protection of their host tissue, CD4^+^ tissue-resident memory cells also contribute to systemic immunity.

## Abbreviations

T_CM_: Central memory T cell
T_EM_: Effector memory T cell
MMR: Measles-mumps-rubella
TCR: T cell receptor
Th: T-helper
PBMC: Peripheral blood mononuclear cell
TT: Tetanus toxoid

## Author Contributions

Conceptualization, J.D. and A.R.; Methodology, C.C., W.J.D. and J.D.; Investigation, C.C., W.J.D., G.A.H. and M.F.M.; Formal Analysis, C.C., W.J.D., P.D., K.T., and J.D.; Resources, T.A., A.S., T.D., A.B., K.S., and HD.C; Interpretation, C.C., W.J.D., HD.V., C.R., K.T., A.R. and J.D.; Writing – Original Draft, J.D.; Writing – Review & Editing, L.S., K.T., A.R., and J.D.; Visualization, C.C., W.J.D, K.T. and J.D.; Supervision, K.T. and J.D.; Project Administration, J.D.; Funding Acquisition, J.D. and A.R. All authors contributed to the final version of the manuscript.

