## Supplemental_Tables for "Mobilization of tissue-resident memory CD4^+^ T lymphocytes and their contribution to a systemic secondary immune reaction"

**Supporting Information Tables**

**Table S1. Key resource table**

| **REAGENT or RESOURCE** | SOURCE | IDENTIFIER |
|---|---|---|
| **Antibodies** | | |
| anti-CD3 APC/Cy7 | Biolegend | Cat#344818; RRID: AB_10645474 |
| anti-CD8 APC/Cy7 | Biolegend | Cat#301016; RRID: AB_314134 |
| anti-CD8 BV785 | Biolegend | Cat#301046; RRID: AB_2563264 |
| anti-CD45 FITC | Biolegend | Cat#304006; RRID: AB_314394 |
| anti-CD45RA BV570 | Biolegend | Cat#304132; RRID: AB_314410 |
| anti-CD56 PE | Biolegend | Cat#318306; RRID: AB_604101 |
| anti-CD69 APC/Cy7 | Biolegend | Cat#310914; RRID: AB_314849 |
| anti-CD154 BV421 | Biolegend | Cat#310824; RRID: AB_2562721 |
| anti-CCR7 A488 | Biolegend | Cat#353206; RRID: AB_10916389 |
| anti-IFNγ PE/Cy7 | Biolegend | Cat#502528; RRID: AB_2123323 |
| anti-IL-2 APC/Cy7 | Biolegend | Cat#500342; RRID: AB_2562855 |
| anti-IL-2 FITC | Biolegend | Cat# 500304; RRID: AB_315091 |
| anti-CD19 BV421 | BD Horizon | Cat# 562440 |
| anti-TNFα APC | BDPharminogen | Cat# 551384 |
| anti-CD4 PE/Cy5.5 | eBioscience | Cat# 35-0047-42; RRID: AB_11218283 |
| anti-KI67 PE | eBioscience | Cat#12-5699-42; RRID: AB_10688373 |
| anti-CD3 FITC | In house | Clone UCHT1 |
| anti-CD4 PE | In house | Clone TT1 |
| anti-CD8 PE/Cy7 | In house | Clone GN11/134D7 |
| anti-CD14 PO | In house | Clone TM1 |
| anti-CD19 PO | In house | Clone BU12 |
| anti-CD3 PerCP | Miltenyi Biotec | Cat# 130-094-965 |
| anti-CD127 PE | Miltenyi Biotec | Cat# 130-109-435 |
| **Chemicals, Peptides, and Recombinant Proteins** | | |
| Buffer EL | Qiagen | Cat# 79217 |
| 5X Lyse/Fix Solution | BD Biosciences | Cat# 558049 |

| | | |
|---|---|---|
| BD FACS Lysing solution | BD Biosciences | Cat# 349202 |
| BD FACS Permeabilizing solution 2 | BD Biosciences | Cat# 347692 |
| Brefeldin A | Biolegend | Cat# 420601 |
| (MMR) live attenuated vaccine-Priorix® | GSK | N/A |
| Qiazol Lysis Reagent | Qiagen | Cat# 79306 |
| X-VIVO 15 Chemically defined medium | Lonza | Cat# BE02-060F |
| CD28 pure - functional grade | Miltenyi Biotec | Cat# 130-093-375 |
| CD40 pure - functional grade | Miltenyi Biotec | Cat# 130-094-133 |
| FcR blocking reagent | Miltenyi Biotec | Cat# 130-059-901 |
| CMV-pp65 | Miltenyi Biotec | Cat# 130-091-824 |
| Measles | Microbix Biosystems | Cat# EL-04-02-001 |
| Mumps | Microbix Biosystems | Cat# El-06-02-001 |
| Rubella | Microbix Biosystems | Cat# EL-05-11-001 |
| Tetanus Toxoid | NIBSC | Cat# 02/232 |
| Q5 High-Fidelity DNA Polymerase | New England Biolabs | Cat# M0491L |
| Uracil-DNA glycosilase (UDG) | New England Biolabs | Cat# M0280L |
| Proleukin (IL-2 clinical use) | Novartis | N/A |
| Human male Ab serum | Sigma-Aldrich | Cat# H4522-100ML |
| CellTrace™ CFSE Cell Proliferation Kit | ThermoFischer | Cat# C34554 |
| Live/Dead™ Fixable Aqua Dead Cell Stain Kit (Pacific Orange) | ThermoFischer | L34957 |
| Penicillin-Streptomycin-Glutamine (100X) | ThermoFischer | 10378016 |
| RPMI Medium 1640 - GlutaMax | ThermoFischer | Cat# 21875-091 |
| RNAse inhibitor | ThermoFischer | Cat# N8080119 |
| **Critical Commercial Assays** | | |
| SMARTer® PCR cDNA synthesis | Clontech | Cat# 634926 |
| TruSeq DNA PCR-Free Library Prep | Clontech | Cat# 20015962 |
| miRNAeasy Micro Kit | Qiagen | Cat# 217084 |
| MinElute PCR Purification Kit | Qiagen | Cat# 28004 |
| QIAquick PCR purification kit | Qiagen | Cat# 28104 |
| PCR-clean-up Gel extraction | MACHEREI-NAGEL | Cat# 740587 |
| Measles virus IgG ELISA | IBL International | Cat# RE57141 |
| Measles virus IgM µ-capture ELISA | IBL International | Cat# RE57151 |
| Mumps virus (Parotitis) IgG ELISA | IBL International | Cat# RE56641 |
| Mumps virus (Parotitis) IgM ELISA | IBL International | Cat# RE56651 |

| | | |
|---|---|---|
| Rubella virus IgG ELISA | IBL International | Cat# RE57081 |
| Rubella virus IgM µ-capture ELISA | IBL International | Cat# RE57091 |
| Tetanus virus IgG ELISA | IBL International | Cat# RE56901 |
| CD154 MicroBead Kit | Miltenyi Biotec | Cat# 130-092-658 |
| Inside stain kit | Miltenyi Biotec | Cat# 130-090-477 |
| **Oligonucleotides** | | |
| **1st strand cDNA synthesis** | | |
| SmartNNNa: template switch adapter with unique molecular identifier (UMI): AAGCAGUGGTAUCAACGCAGAGU NNNN U NNNN U NNNNN UCTT gggg | Metabion | (Mamedov et al., 2013)(1) |
| bc1R: primer for cDNA synthesis, human TCR beta mRNA CAG TAT CTG GAG TCA TTG A | Metabion | (Mamedov et al., 2013) |
| **1st PCR amplification** | | |
| bc2R: nested primer 1, human TCR beta library TGC TTC TGA TGG CTC AAA CAC | Metabion | (Mamedov et al., 2013) |
| Step-out primers 1 with sample barcodes. Anneals on the template switch adapter | | (Mamedov et al., 2013) |
| Na-SB2-M1: CGA GCG TGA CGA CGA CAG TAG TCG TGG TAT CAA CGC AGA GT | Metabion | (Mamedov et al., 2013) |
| Na-SB4-M1: CGA GCG TGA CGA CGA CAG TCA TCG TGG TAT CAA CGC AGA GT | Metabion | (Mamedov et al., 2013) |
| Na-SB5-M1: CGA GCG TGA CGA CGA CAG GAT TCG TGG TAT CAA CGC AGA GT | Metabion | (Mamedov et al., 2013) |
| Na-SB6-M1: CGA GCG TGA CGA CGA CAG GTC TTG TGG TAT CAA CGC AGA GT | Metabion | (Mamedov et al., 2013) |
| Na-SB7-M1: CGA GCG TGA CGA CGA CAG AGT CTG TGG TAT CAA CGC AGA GT | Metabion | (Mamedov et al., 2013) |
| Na-SB8-M1: CGA GCG TGA CGA CGA CAG ACT TCA GTG GTA TCA ACG CAG AGT | Metabion | (Mamedov et al., 2013) |
| Na-SB9-M1: CGA GCG TGA CGA CGA CAG ATC CTA GTG GTA TCA ACG CAG AGT | Metabion | (Mamedov et al., 2013) |
| Na-SB10-M1: CGA GCG TGA CGA CGA CAG CAA CTT GTG GTA TCA ACG CAG AGT | Metabion | (Mamedov et al., 2013) |

| | | |
|---|---|---|
| Na-SB11-M1:<br>CGA GCG TGA CGA CGA CAG CCT AAT GTG GTA TCA ACG CAG AGT | Metabion | (Mamedov et al., 2013) |
| Na-SB12-M1:<br>CGA GCG TGA CGA CGA CAG CGG TCT GTG GTA TCA ACG CAG AGT | Metabion | (Mamedov et al., 2013) |
| Na-SB13-M1:<br>CGA GCG TGA CGA CGA CAG CTC GGT GTG GTA TCA ACG CAG AGT | Metabion | (Mamedov et al., 2013) |
| Na-SB14-M1:<br>CGA GCG TGA CGA CGA CAG GCT CTG GTG GTA TCA ACG CAG AGT | Metabion | (Mamedov et al., 2013) |
| Na-SB16-M1:<br>CGA GCG TGA CGA CGA CAG TAA TCC GTG GTA TCA ACG CAG AGT | Metabion | (Mamedov et al., 2013) |
| Na-SB17-M1:<br>CGA GCG TGA CGA CGA CAG TCT GGC GTG GTA TCA ACG CAG AGT | Metabion | (Mamedov et al., 2013) |
| Na-SB18-M1:<br>CGA GCG TGA CGA CGA CAG TGG CTC GTG GTA TCA ACG CAG AGT | Metabion | (Mamedov et al., 2013) |
| Na-SB19-M1:<br>CGA GCG TGA CGA CGA CAG TTC AAC GTG GTA TCA ACG CAG AGT | Metabion | (Mamedov et al., 2013) |
| Na-SB20-M1:<br>CGA GCG TGA CGA CGA CAG GTC TCG GTG GTA TCA ACG CAG AGT | Metabion | (Mamedov et al., 2013) |
| **2nd PCR amplification** | | |
| Nested primer 2, human TCR beta. Anneals to the switch adapter | | |
| N2Na:<br>NNC GAG CGT GAC GAC GAC AG | Metabion | (Mamedov et al., 2013) |
| N3Na:<br>NNN CGA GCG TGA CGA CGA CAG | Metabion | (Mamedov et al., 2013) |
| N4Na:<br>NNN NCG AGC GTG ACG ACG ACA G | Metabion | (Mamedov et al., 2013) |
| Reverse primer for TCR beta chain with sample barcodes | Metabion | (Mamedov et al., 2013) |
| bc3R-SB2r:<br>TAG TCA CAC STT KTT CAG GTC CTC | Metabion | (Mamedov et al., 2013) |
| bc3R-SB4r:<br>TCA TCA CAC STT KTT CAG GTC CTC | Metabion | (Mamedov et al., 2013) |

| | | |
|---|---|---|
| bc3R-SB5r:<br>GAT TCA CAC STT KTT CAG GTC CTC | Metabion | (Mamedov et al., 2013) |
| bc3R-SB6r:<br>GTC TTA CAC STT KTT CAG GTC CTC | Metabion | (Mamedov et al., 2013) |
| bc3R-SB7r:<br>AGT CTA CAC STT KTT CAG GTC CTC | Metabion | (Mamedov et al., 2013) |
| bc3R-SB8r:<br>ACT TCA ACA CST TKT TCA GGT CCT C | Metabion | (Mamedov et al., 2013) |
| bc3R-SB9r:<br>ATC CTA ACA CST TKT TCA GGT CCT C | Metabion | (Mamedov et al., 2013) |
| bc3R-SB10r:<br>CAA CTT ACA CST TKT TCA GGT CCT C | Metabion | (Mamedov et al., 2013) |
| bc3R-SB11r:<br>CCT AAT ACA CST TKT TCA GGT CCT C | Metabion | (Mamedov et al., 2013) |
| bc3R-SB12r:<br>CGG TCT ACA CST TKT TCA GGT CCT C | Metabion | (Mamedov et al., 2013) |
| bc3R-SB13r:<br>CTC GGT ACA CST TKT TCA GGT CCT C | Metabion | (Mamedov et al., 2013) |
| bc3R-SB14r:<br>GCT CTG ACA CST TKT TCA GGT CCT C | Metabion | (Mamedov et al., 2013) |
| bc3R-SB16r:<br>TAA TCC ACA CST TKT TCA GGT CCT C | Metabion | (Mamedov et al., 2013) |
| bc3R-SB17r:<br>TCT GGC ACA CST TKT TCA GGT CCT C | Metabion | (Mamedov et al., 2013) |
| bc3R-SB18r:<br>TGG CTC ACA CST TKT TCA GGT CCT C | Metabion | (Mamedov et al., 2013) |
| bc3R-SB19r:<br>TTC AAC ACA CST TKT TCA GGT CCT C | Metabion | (Mamedov et al., 2013) |
| bc3R-SB20r:<br>GTC TCG ACA CST TKT TCA GGT CCT C | Metabion | (Mamedov et al., 2013) |
| **Software and Algorithms** | | |

| | |
|---|---|
| BD FACSDiva 6 | BD Biosciences |
| MACSQuantify™ software | Miltenyi Biotec |
| Flowjo 9.9.4 / 10.2 | Tree Star |
| GraphPad Prism 5.x | GraphPad Software |

**Table S2. Study subjects**

| Donor | Age (years) | Gender | Antibody titers pre-vaccination | | |
|---|---|---|---|---|---|
| | | | Measles | Mumps | Rubella |
| | | | (mIU/mL) | (U/mL) | (IU/mL) |
| | | | Cut off: 220 | Cut off: 350 | Cut off: 10) |
| MMR1 | 28 | F | 400 | 700 | 161 |
| MMR2 | 34 | F | 3300 | 2200 | 42 |
| MMR3 | 32 | F | 1400 | 2600 | 44 |
| MMR4 | 29 | F | -ve | 2500 | 424 |
| MMR5 | 26 | F | 1900 | 860 | 44 |
| MMr6 | 43 | F | 9200 | 2500 | 95 |
| MMR7 | 42 | M | 11000 | 4400 | 26 |
| MMR8 | 31 | M | 920 | 3800 | 187 |
| MMR9 | 38 | M | 360 | 350 | 16 |
| MMR10 | 37 | M | 4200 | 4600 | 107 |
| MMR11 | 34 | F | +ve | +ve | +ve |
| Ctrl1 | 57 | M | +ve | +ve | 237.50 |
| Ctrl2 | 34 | F | +ve | +ve | 127.50 |
| Ctrl3 | 24 | F | +ve | +ve | -ve |
| Ctrl4 | 31 | F | +ve | +ve | 440.5 |

Information about age, gender, and measles, mumps, and rubella virus antigen-specific antibody titers of study subjects pre-vaccination. –ve, negative; +ve, positive.

**Table S3. Sequencing metadata for TCR CDR3 Vβ repertoires.**

| Sample | Replicates (N° cells) | | Mapped reads | | UMI counts | | Clonotypes | |
|---|---|---|---|---|---|---|---|---|
| | a | b | a | b | a | b | a | b |
| MMR10_d0 | 2500 | 2500 | 749116 | 266776 | 175 | 269 | 106 | 150 |
| MMR10_d1 | 2500 | 2500 | 651958 | 242883 | 638 | 523 | 360 | 282 |
| MMR10_d14 | 2500 | 2500 | 284930 | 367200 | 783 | 763 | 395 | 393 |
| MMR11_d0 | 2500 | 2500 | 356605 | 194558 | 376 | 399 | 302 | 314 |
| MMR11_d1 | 2500 | 2500 | 587611 | 263078 | 773 | 835 | 558 | 602 |
| MMR11_d14 | 2500 | 2500 | 849199 | 300429 | 404 | 463 | 306 | 343 |

PBMCs from vaccinees MMR10 and MMR11 stimulated with measles and two replicates (a and b) and from each 2,500 antigen-reactive $CD154^+CD69^+$ memory $CD4^+$ T cells were isolated before (d0) and 1 (d1) and 14 (d14) days after MMR vaccination. Total RNA was extracted from each set of cells and subjected to parallel library preparation. The mapped reads, the counts of unique molecular identifier (UMI; tagging the quantity of unique mRNA transcripts), CDR3 clonotypes from both replicates are shown.
