## Supplemental Figures for "Mobilization of tissue-resident memory CD4^+^ T lymphocytes and their contribution to a systemic secondary immune reaction"

### Supporting Information Figures

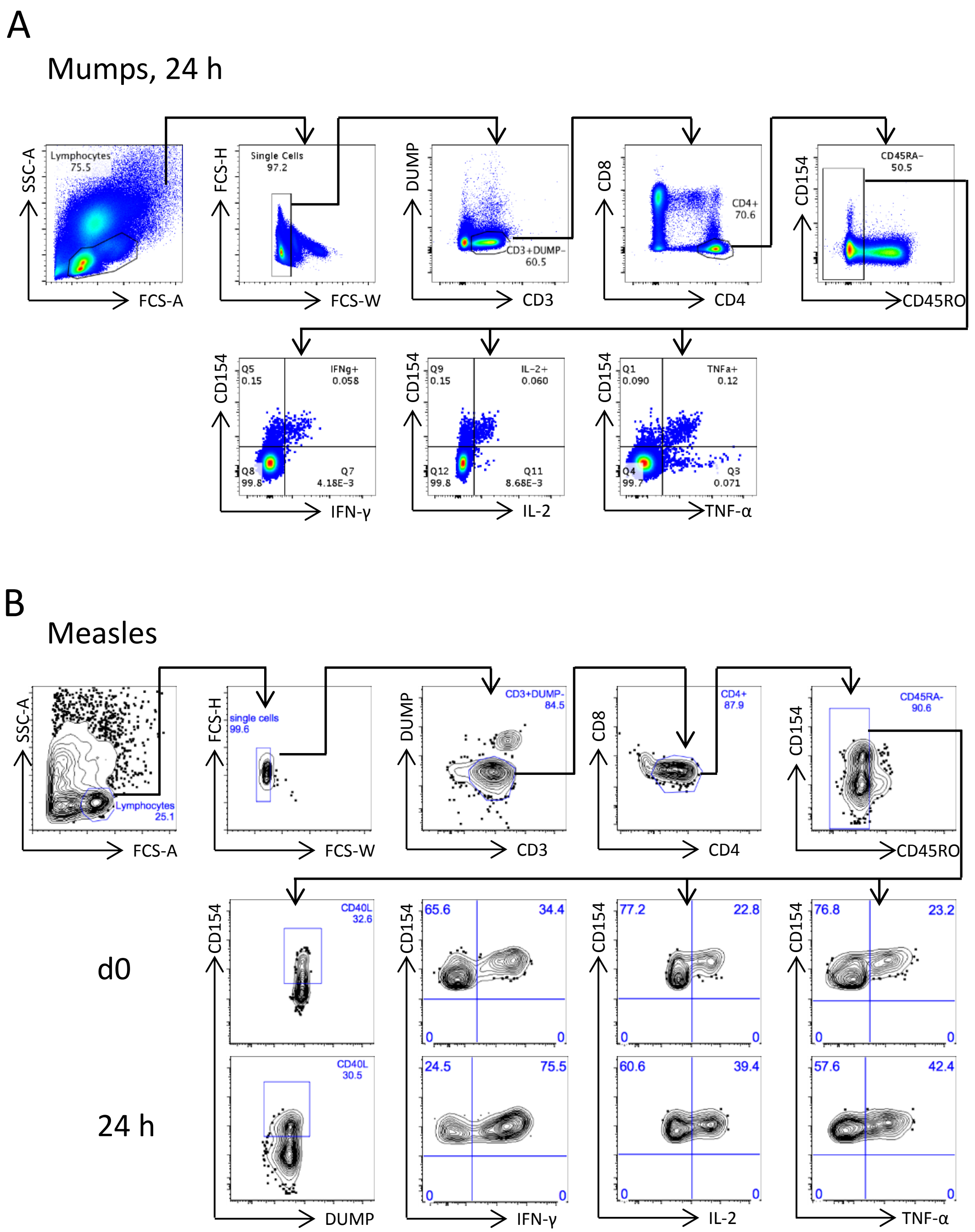


**Figure S1. Gating strategy for analyzing antigen-specific memory CD4$^+$ T cell responses in previously naïve and immune donors before and post-MMR vaccination**. PBMCs from vaccinees were stimulated with the indicated vaccine antigens, and the induced cytokine production (IL-2, TNF-α, and IFN-γ) in memory CD4$^+$ T cells was examined according to CD154 expression by conventional intracellular cytokine staining (A) or by CD154 pre-enrichment (B) at the indicated time points.

A

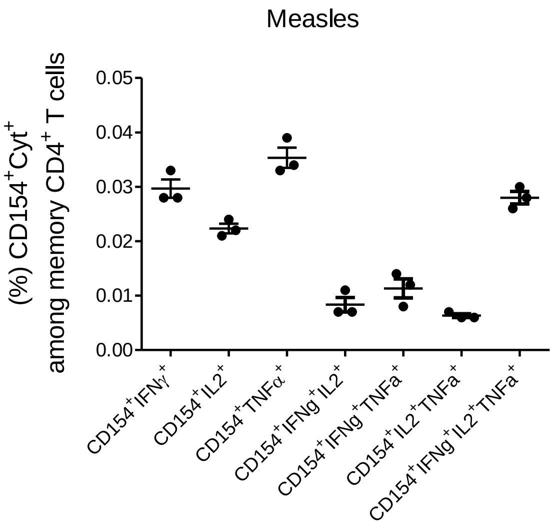


B

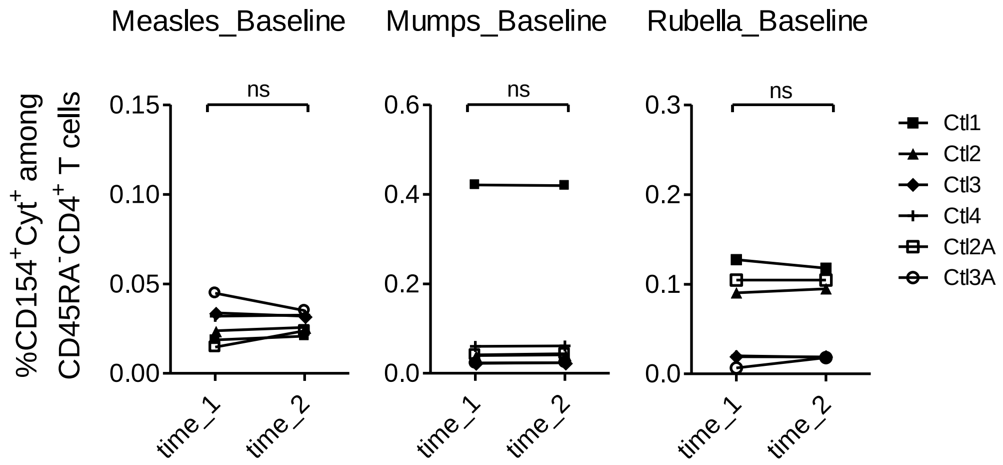


**Figure S2. Assay control for analyzing antigen-specific memory CD4⁺ T cell responses**. PBMCs from baseline control donors (i.e. without MMR vaccination) were stimulated with the indicated vaccine antigens, and the induced cytokine production (IL-2, TNF-α, and IFN-γ) in memory CD4⁺ T cells was examined according to CD154 expression by intracellular cytokine staining. Frequencies of antigen-reactive CD154⁺ memory CD4⁺ T cells producing either and/or of the analyzed cytokines from technical replicates (A) or from two consecutive days between time points 1 and 2 (B) are shown.

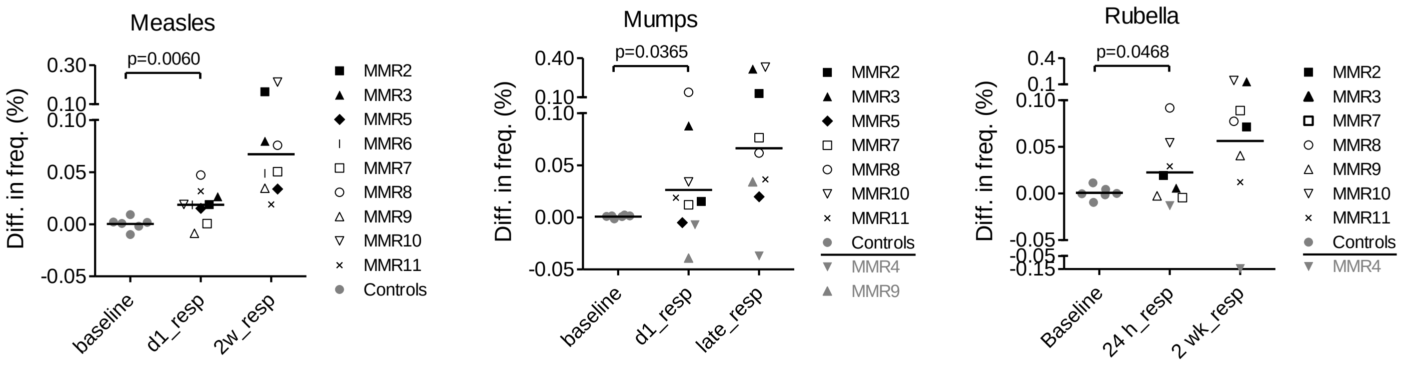


**Figure S3. Quality control of antigen-specific memory CD4⁺ T cell responses after MMR vaccination**. PBMCs from vaccinees were stimulated with the indicated vaccine antigens, and the induced cytokine production (IL-2, TNF-α, and IFN-γ) in memory CD4⁺ T cells was examined according to CD154 expression by intracellular cytokine staining. Differences in frequencies of antigen-reactive CD154⁺ memory CD4⁺ T cells producing either and/or of the analyzed cytokines from two consecutive days between time points 1 and 2 without vaccination (i.e. baseline; Figure S2B) and from two consecutive days before and 1 day after vaccination from vaccinees with 2-wk immune responses were compared. Vaccinees that do not generate 2 wk immune responses were labeled in grey and excluded for statistics treatment. Unpaired t-test with Welch's correction was performed and one-tailed *P* value is shown.

A

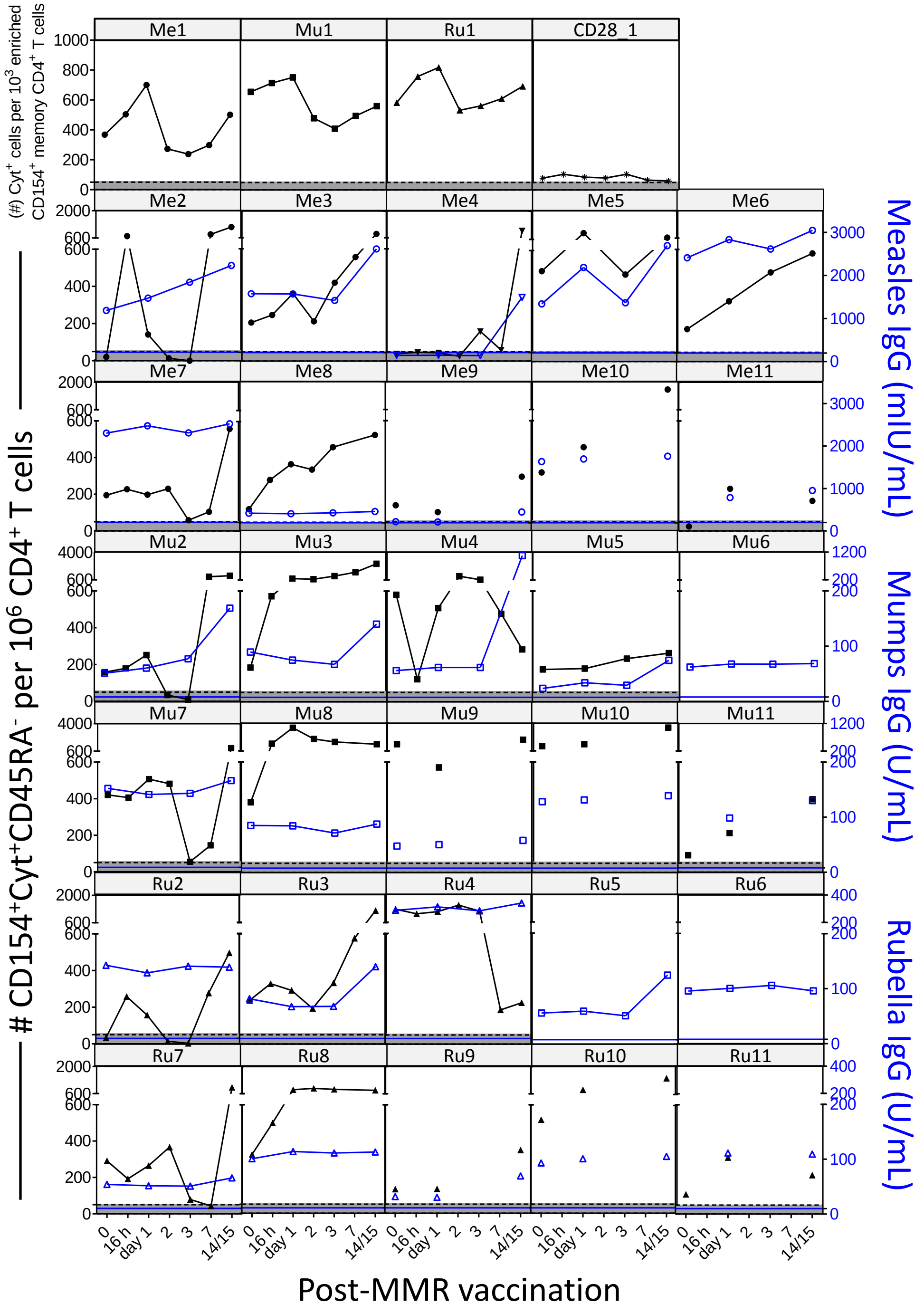

Me1
Mu1
Ru1
CD28_1
(#) Cyt+ cells per 10^3 enriched CD154+ memory CD4+ T cells
Me2
Me3
Me4
Me5
Me6
Me7
Me8
Me9
Me10
Me11
Mu2
Mu3
Mu4
Mu5
Mu6
Mu7
Mu8
Mu9
Mu10
Mu11
Ru2
Ru3
Ru4
Ru5
Ru6
Ru7
Ru8
Ru9
Ru10
Ru11
### CD154+Cyt+CD45RA- per 10^6 CD4+ T cells
Measles IgG (mIU/mL)
Mumps IgG (U/mL)
Rubella IgG (U/mL)
0
16 h
day 1
2
3
7
14/15
Post-MMR vaccination

B

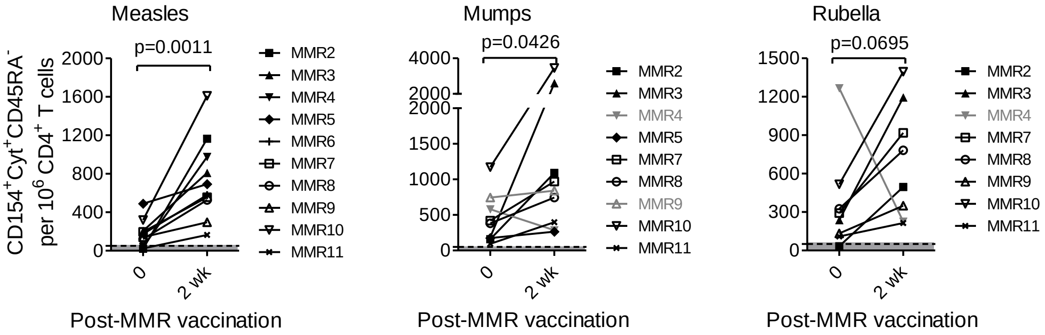


C

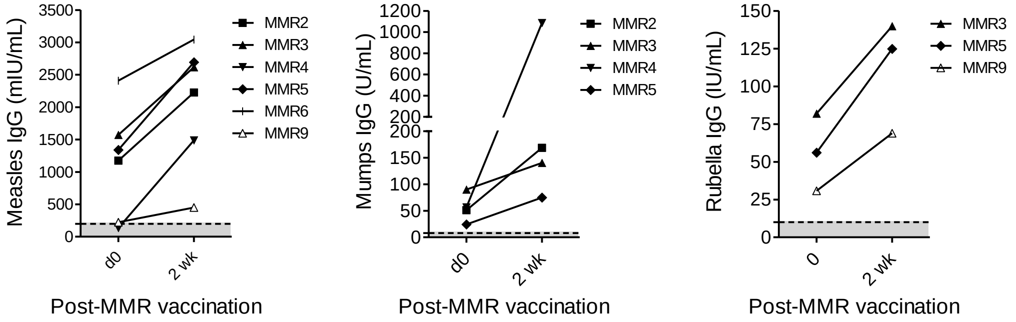


**Figure S4. Antigen-specific memory CD4⁺ T cell and antibody responses in naïve and previously immune donors before and post-MMR vaccination**. (A, B) PBMCs from vaccinees were stimulated with measles, mumps, or rubella and analyzed for antigen-reactive CD154⁺cytokine⁺ (black line or dot) memory T cells (memory CD4⁺ T cells expressing CD154 (CD40L) and one or more of the cytokines IL-2, TNF-α, and IFN-γ) per $10^6$ CD4⁺ T cells or per $10^3$ enriched CD154⁺ memory CD4⁺ T cells as referenced to the left y-axis at indicated time points before and after vaccination (A) and comparisons of antigen-reactive cells of before and 2 wk after vaccination (B). Serum from vaccinees analyzed for measles-, mumps-, or rubella-specific IgG antibody titers (blue line or dot) at time points before and after MMR vaccination as referenced to the right y-axis (A). In B, data shown in grey indicate less than 30% of responses detected at 2 wk after vaccination as compared to d0 without vaccination. (C) Vaccinees anaylzed displaying more than 30% increase in measles-, mumps-, or rubella-specific IgG antibody titers. The dotted black or blue line indicates the minimum threshold for detection with the respective assay and the grey areas underneath values below reliable detection limits.

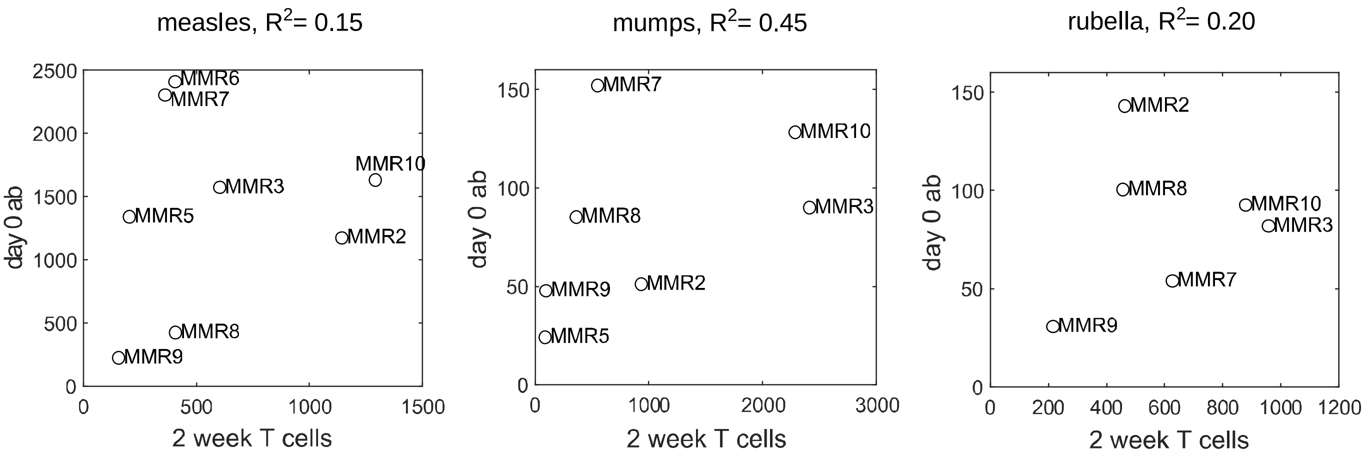


**Figure S5. Memory T cell and responses 2 weeks after MMR vaccination and their related antibody titers before vaccination** Coefficient of determination ($R^2$) was calculated based on antigen specific antibody titer before vaccination (d0) and the increased absolute numbers of measles-, mumps-, or rubella-reactive memory T cells per $10^6$ $CD4^+$ T cells between d0 before and d14 after MMR vaccination.

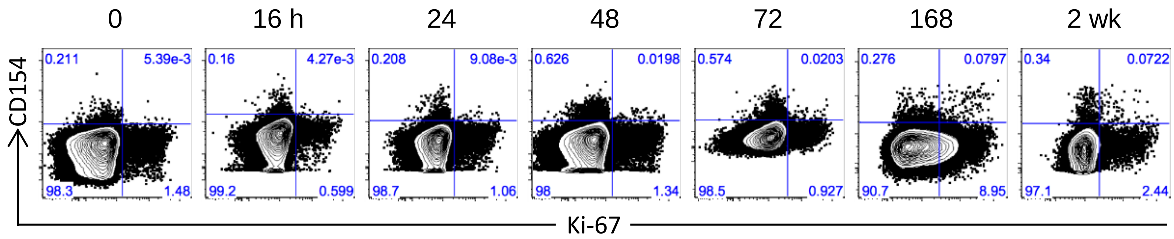


**Figure 6. Kinetic expression of Ki-67 on antigen-specific CD154$^+$ memory CD4$^+$ T cells in previously immune donors before and post-MMR vaccination**. PBMCs from vaccinees were stimulated with measles, mumps, or rubella, and the Ki-67 expression on memory CD4$^+$ T cells was examined according to CD154 expression by conventional intracellular staining. Data shown are representative analysis of Ki-67 expression on mumps-reactive CD154$^+$ memory CD4$^+$ T cells from vaccinee MMR3.

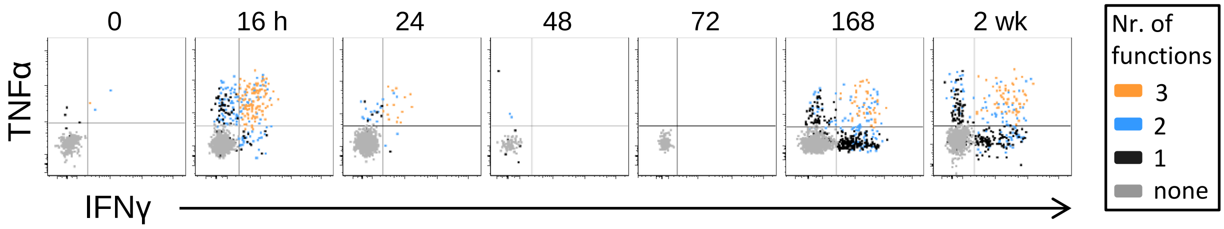


**Figure S7. Kinetics of cytokine production on antigen-specific CD154⁺ memory CD4⁺ T cells in previously immune donors before and post-MMR vaccination**. PBMCs from vaccinees were stimulated with measles, mumps, or rubella, and the and the induced cytokine production (IL-2, TNF-α, and IFN-γ) in memory $CD4^+$ T cells was examined according to CD154 expression by conventional intracellular cytokine staining. Numbers of functions, cells producing 1 (black dots), 2 (blue dots), 3 (orange dots), or none (grey dots) of the three analyzed cytokines are shown. Data shown are representative analysis of measles-reactive $CD154^+$ memory $CD4^+$ T cells from vaccinee MMR2.

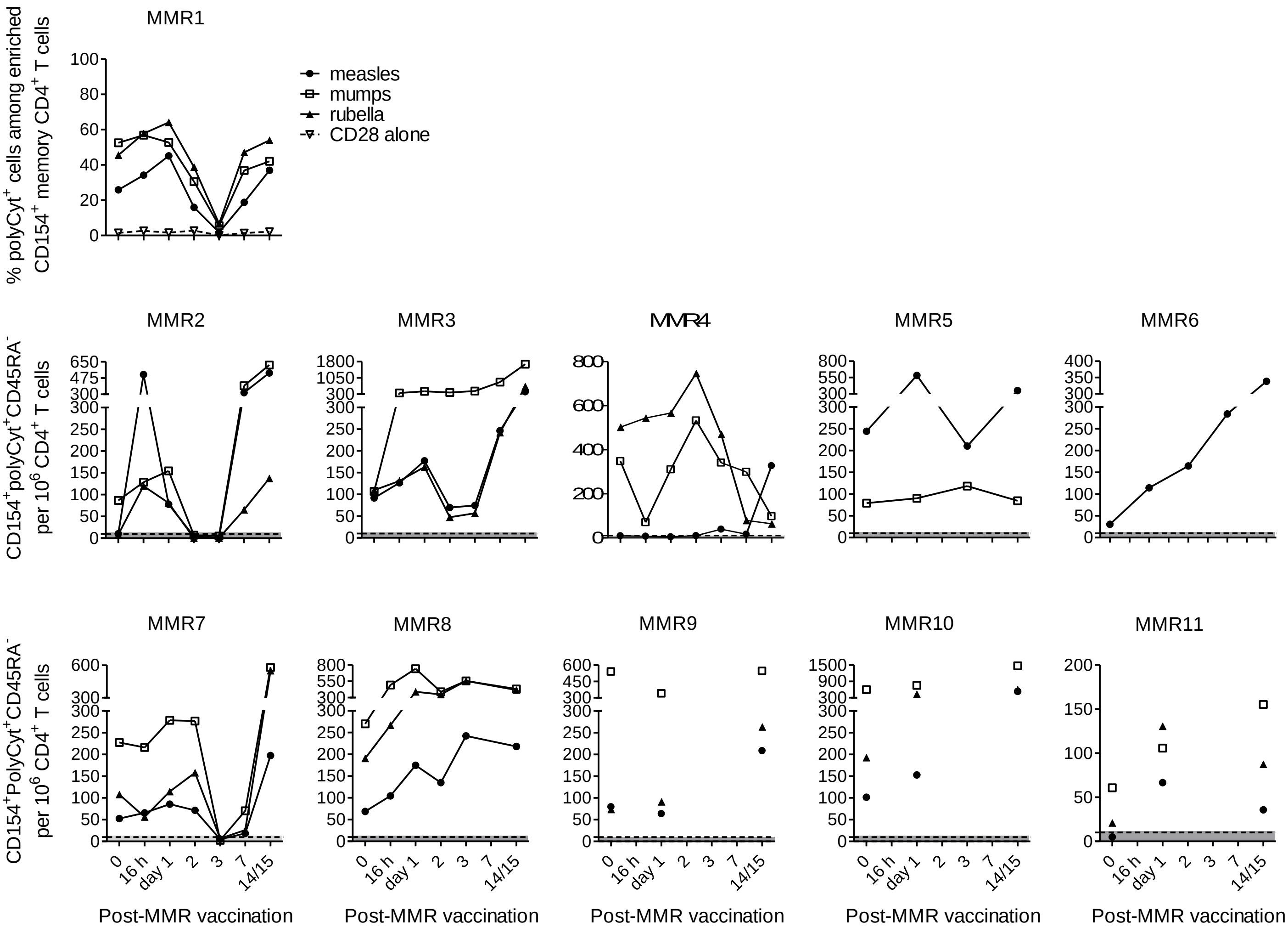


**Figure S8. Antigen-specific polyfunctional CD4⁺ T cell and antibody responses in naïve and previously immune donors before and post-MMR vaccination** PBMCs from vaccinees were stimulated with measles, mumps, or rubella and analyzed for memory T cells expressing CD154 and two or three of the cytokines IL-2, IFN-γ or TNF-α, i.e. antigen-reactive CD154⁺ poly-cytokine⁺ memory T cells per $10^6$ CD4⁺ T cells. The dotted line indicates the minimum threshold for detection with the respective assay and the grey areas underneath values below reliable detection limits.

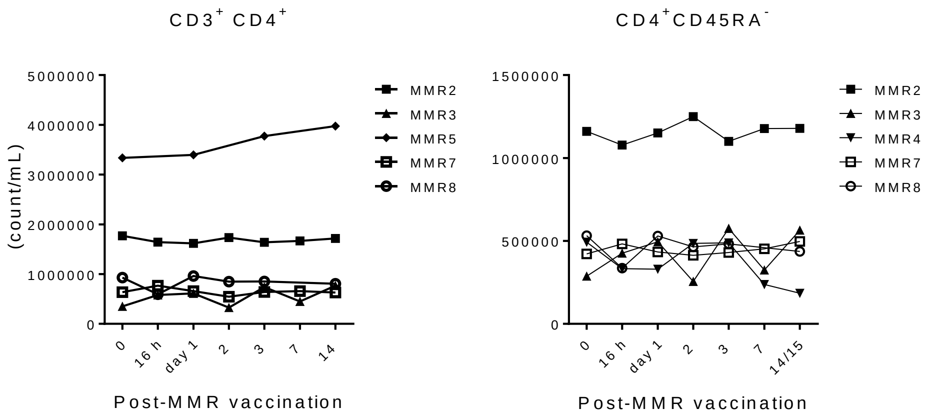


**Figure S9. $CD3^+CD4^+$ T cell and $CD4^+CD45RA^-$ memory T cell counts per mL blood at various time points pre- and post-MMR vaccination** Blood samples from vaccinees were analyzed for $CD3^+CD4^+$ T cell counts. Based on the frequency of $CD45RA^-$ among $CD4^+$ cells, $CD4^+CD45RA^-$ memory T cell counts were calculated.

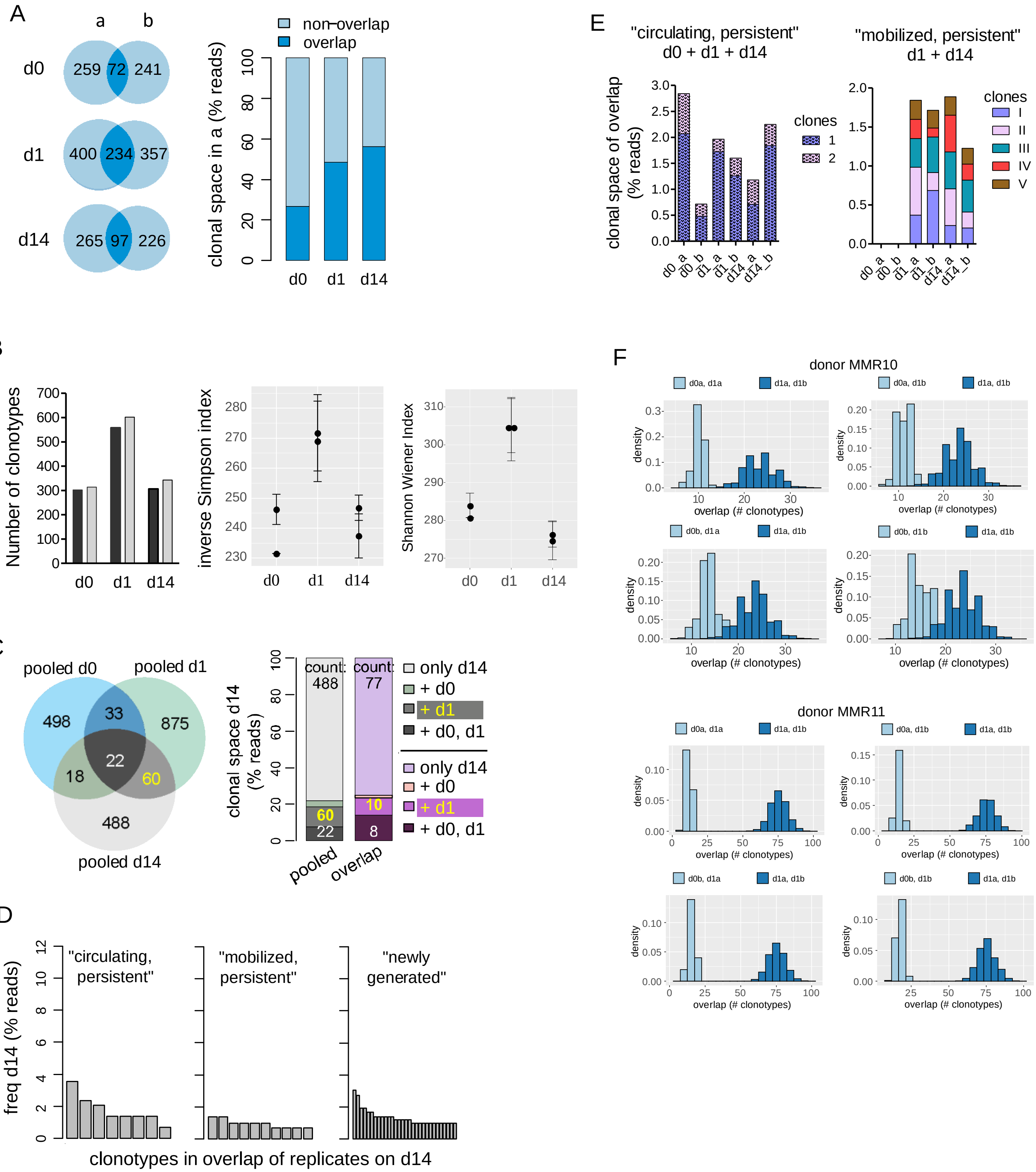
A
a
b
d0
259
72
241
d1
400
234
357
d14
265
97
226
non-overlap
overlap
clonal space in a (% reads)
0
20
40
60
80
100
d0
d1
d14
B
Number of clonotypes
0
100
200
300
400
500
600
700
d0
d1
d14
inverse Simpson index
230
240
250
260
270
280
Shannon Wiener Index
270
280
290
300
310
C
pooled d0
pooled d1
pooled d14
498
33
875
22
18
60
488
clonal space d14 (% reads)
count: 488
count: 77
60
22
10
8
pooled
overlap
only d14
+ d0
+ d1
+ d0, d1
only d14
+ d0
+ d1
+ d0, d1
D
freq d14 (% reads)
0
2
4
6
8
10
12
"circulating, persistent"
"mobilized, persistent"
"newly generated"
clonotypes in overlap of replicates on d14
E
"circulating, persistent"
d0 + d1 + d14
"mobilized, persistent"
d1 + d14
clonal space of overlap (% reads)
clones
1
2
clones
I
II
III
IV
V
d0_a
d0_b
d1_a
d1_b
d14_a
d14_b
F
donor MMR10
d0a, d1a
d1a, d1b
d0a, d1b
d0b, d1a
d0b, d1b
density
overlap (# clonotypes)
donor MMR11

**Figure S10. Supporting information for TCR repertoire analysis in Figure 4. TCR CDR3 Vβ clonotypes analysis of measles-reactive $CD4^+$ memory T cells after MMR vaccination** PBMCs were isolated before (d0) and 1 (d1) and 14 (d14) days after MMR vaccination from vaccinees MMR10 and MMR11, and restimulated *ex vivo* with measles for measles-reactive $CD154^+CD69^+$ memory $CD4^+$ T cells. In A-E, data shown for MMR11 (A) Analysis of two replicates at each time point, shown is the “overlap”, i.e. the amount of clonotypes detected in both replicates, in terms of clonotype count (Venn diagrams) and frequency (bar graphs) (B) Diversity of the TCR repertoire. Shown are the number of clonotypes, the inverse Simpson index and the Shannon Wiener Index at three time points, for both replicates in A. Error bars indicate standard deviation after repetitive down-sampling. (C) Venn diagrams show comparative numbers of clonotypes in “pooled” repertoires, i.e. clonotypes detected in at least one replicate. Clonal space (bar graph) corresponds to the fraction of reads detected on d14 “overlap” or “pooled” that were additionally present in at least one of the replicates at the indicated time points. Numbers in the bar plot indicate numbers of clonotypes (for pooled replicates, these numbers match the Venn diagram). (D) Frequency distributions of “circulating, persistent” (d14+d0+d1), “mobilized, persistent” (d14+d1) and “newly generated (only d14) clonotypes (see C), with respect to the overlap of replicates on d14. For “newly generated”, only the 30 most frequent clonotypes are shown. (E) Contribution of each replicate to the “circulating, persistent” and “mobilized, persistent” clonotypes. Clonal space is given as percentage of reads in the overlap of replicates on indicated days. Only clonotypes present in all applicable samples are shown. (F) Comparison of the overlap of samples taken from both vaccinees MMR10 and MMR11 on different days or taken on the same day. Shown are the distributions of indicated sample overlaps after repetitive down-sampling to the number of reads in the smallest among all samples taken on d0 and d1.
